# Building evolutionary resilience: a framework for managing safe haven populations

**DOI:** 10.1101/2025.08.11.669227

**Authors:** Alex Slavenko, Collin W Ahrens, Adrian D Manning, Andrew R Weeks

## Abstract

Safe havens are widely used to mitigate the impacts of invasive predators on threatened species, yet their isolation often disrupts evolutionary processes and can leave them vulnerable to environmental change and stochastic events. We present a framework for integrating evolutionary principles into safe haven management, termed 500-in-5, which establishes a metapopulation of >500 breeding individuals distributed across five geographically separated safe havens spanning environmental gradients. Using genetic and demographic simulations for two threatened Australian mammals, we demonstrate how this approach can enhance adaptive potential and persistence. Spatially explicit population models predict that many existing safe havens are at risk of collapse within decades. However, genetic simulations show that selection on standing genetic variation can drive fitness gains within tens of generations, improving population resilience to novel conditions. Integrating these processes reveals that safe haven networks can promote adaptation and resilience, substantially improving long-term persistence and providing robust sources for reintroductions.

## Introduction

The accelerating biodiversity crisis demands urgent, innovative conservation strategies to protect threatened species. In many regions, safe havens (alternatively known as sanctuaries or feral predator-free fenced reserves and islands) play a pivotal role in preventing extinctions, particularly in landscapes dominated by invasive predators (*e.g.*, in Australia; Hayward *et al*. 2014; Legge *et al*. 2018). Safe havens are designed to secure populations through initial intensive predator control and hands-on management, often enabling species to persist where they would otherwise be lost (Innes *et al*. 2015).

However, safe haven populations face inherent evolutionary challenges. Often established from a limited number of founders, with small population sizes and lacking natural dispersal, many suffer from inbreeding, genetic drift, and reduced adaptive potential (Willi *et al*. 2006; Weeks *et al*. 2011). These issues constrain genetic diversity and limit their ability to respond to environmental change, making them vulnerable to extinction (Hayward & Kerley 2009).

Building evolutionary resilience, defined as the capacity of populations to adapt and persist under environmental change (Hoffmann & Sgrò 2011; Sgrò *et al*. 2011), is therefore critical for long-term conservation outcomes (Evans *et al*. 2022).

One pathway to enhance evolutionary resilience is to replicate natural evolutionary processes within safe haven networks. This involves maintaining multiple, demographically stable populations distributed across habitats with varying environmental pressures, including some sites that expose populations to “stressors” (*e.g.*, warmer or drier climates, increased predation risk) and others that provide more benign “core” conditions. This approach contrasts with current practices, which often prioritise controlled, relatively stress-free environments, with species typically confined to a small number of sites, often lacking coordinated management to maintain genetic diversity or to apply selection pressures that could enhance adaptive potential (Mace & Purvis 2008).

Standing genetic variation (SGV), the pool of alleles already present in a population (Barrett & Schluter 2008), is critical for rapid adaptation to changing conditions. In small populations, novel beneficial mutations are rare (Whitlock & Bürger 2009), but existing SGV can enable selection to drive adaptive change when environmental conditions shift (Lai *et al*. 2019).

Establishing multiple safe haven populations from a common heterogeneous genetic source and exposing them to distinct environmental stressors allows natural selection to act differently across sites, generating adaptive variation unlikely to arise in a single, benign population. Importantly, maintaining some gene flow among sites mimics dispersal and increases within-safe haven genetic variation across the metapopulation (Whitlock & Bürger 2009; Orr 2010), further enhancing adaptive potential.

Here we propose the “500-in-5” framework, which integrates evolutionary principles into safe haven management by establishing at least 500 breeding individuals distributed across five geographically distinct safe havens. The threshold of >500 breeding individuals is based on population genetic and evolutionary models, which indicate the minimum population size required to maintain long-term evolutionary potential and minimise the negative effects of genetic drift (Franklin 1980; Lande 1995; Franklin & Frankham 1998; Willi *et al*. 2006; Weeks *et al*. 2011; Frankham *et al*. 2014). The five safe havens enable a metapopulation approach across benign (“core”) and stressor (“stressor”) conditions to promote adaptive evolution.

Periodic translocations between sites will mimic natural dispersal, facilitating gene flow and spreading adaptive alleles. We use combine spatially explicit population modelling and genetic simulations to explore the application of the 500-in-5 framework, using two threatened Australian mammals as case studies and climate change as the stress.

## Methods

### Modelling population trajectories under climate change

To demonstrate how sites may be selected for “core” and “stressor” safe havens for a 500-in-5 program, we explore a case study focussing on two threatened Australian marsupials, the eastern barred bandicoot (*Perameles gunnii*) and the eastern quoll (*Dasyurus viverrinus*).

These species were selected because both:

a) Have similar historical distributions in southeastern Australia (Fig. S1).
b) Were extirpated from the mainland following European colonisation due to introduced predators (foxes and cats), with remnant wild populations only in Tasmania (Jones & Rose 2001; Kinnear *et al*. 2002; DSE Victoria 2009).
c) Have been successfully re-introduced to the mainland in predator-free safe havens.

Climate change was chosen as a stressor, since southeastern Australia is predicted to experience rapid climatic changes during the 21^st^ century leading to warmer, drier conditions (Grose *et al*. 2015).

We implemented a spatially explicit population modelling (SEPM) framework in R v4.5.0 (R Core Team 2025), adapted from Tomlinson *et al*. (2023), to simulate landscape-level population processes integrated with climate suitability models. Briefly, we used current and historical distribution records of the two species to build climate envelope models (Nogués-Bravo 2009), which we used to project past, present, and future climatic suitability across their ranges. We coupled these suitability trends with process-based population models to simulate survival, fecundity, density-dependent population growth, and o@ake (Fordham *et al*. 2021).

We estimated population parameters using external validation targets, sampling parameters from simulations that most closely matched each species’ extirpation history. These parameters were used to simulate future population trajectories in existing and potential safe havens across Australia to predict climate change impacts on safe haven populations (*e.g.*, persist, decline or collapse).

The full methodology behind the simulation approach is in Supporting Information S1.

### Genetic simulations

We hypothesised that establishing safe haven populations across contrasting climatic conditions would elicit adaptive responses. To explore this, we used the 500-in-5 framework and simulated five populations in a Wright-Fisher model with SliM v4.3 (Haller & Messer 2023) implemented in SLiMgui. We first simulated a single large population of 1,000 individuals over 5,000 generations without selection, allowing neutral mutations (1x10^-8^) to accumulate across ten “genes” (genomic regions). This generated a shared background of SGV.

At generation 5,001, five new populations were established by randomly sampling 100 individuals, representing five safe havens with similar starting genetic backgrounds. After the split, fitness effects were applied to five of the ten regions (the remaining five remained neutral) to mimic polygenic adaptation to climate (Barghi *et al*. 2020). The fitness contribution of each adaptive allele was scaled with allele frequency by summing each allele frequency divided by 1,000, ensuring very small per-variant effects.

To represent environmental heterogeneity, two populations were simulated under benign conditions (baseline fitness multiplier = 1; “core” sites), two under mild stress (fitness multiplier = 0.1; mild “stressor” sites), and one under high stress (fitness multiplier = 0.01; strong “stressor” site). These values were chosen to approximate the gradient of selective pressures expected under differing climatic conditions: from relatively stable habitats where fitness is primarily constrained by demographic stochasticity, to increasingly stressful environments where selection acts more strongly on adaptive traits. Simulations continued for 1,000 generations, recording allele frequencies for all adaptive and neutral loci at each generation. We then used linear models to estimate the rate of fitness change per generation for each population.

### Spatial and adaptive modelling

We used the same SEPM framework applied to identify potential “core” and “stressor” sites to examine how changes in fitness could influence safe haven population trajectories. We simulated three scenarios representing differing levels of adaptive response to changing climatic conditions—low, moderate, and high fitness increases—reflecting the intended outcome of implementing the 500-in-5 program across a climatic gradient.

For each cell and year, we incrementally increased simulated climatic suitability by 0.001 (low), 0.002 (moderate) or 0.003 (high) multiplied by the number of years since the start of the simulation, up to a maximum climatic suitability of 1. These correspond to annual fitness increases of 0.1%, 0.2% and 0.3%, or cumulative fitness increase of 7.6%, 15.2% and 22.8%, respectively, by the end of the century.

Simulations were run for existing safe haven sites containing established populations of *P. gunnii* and *D. viverrinus*. For each site and fitness scenario, projected population trajectories were compared to a baseline scenario with no change in fitness (represented by simulations under the “overlapping niches” scenario; see Supporting Information S1). This allowed us to evaluate how adaptive responses to novel climatic conditions could alter long-term population trajectories in sanctuary populations.

## Results

### Modelling sanctuary sites

Our SEPM simulations projected that *P. gunnii* and *D. viverrinus* will decline throughout the 21^st^ century across most of their historical ranges, even if reintroduced into safe havens (Fig. 1). Many existing safe haven populations were also projected to decline (Figs. 2 & S14–15): for *P. gunnii*, all were predicted to collapse by ∼2080, apart from Tiverton and Hamilton Community Parklands (Fig. 2A); for *D. viverrinus*, most populations persisted to 2100 but declined to very small sizes approaching collapse in the latter half of the century (Fig. 2B). The exception was Jervis Bay, which is likely to go extinct much earlier (∼2040).

**Figure 1.**
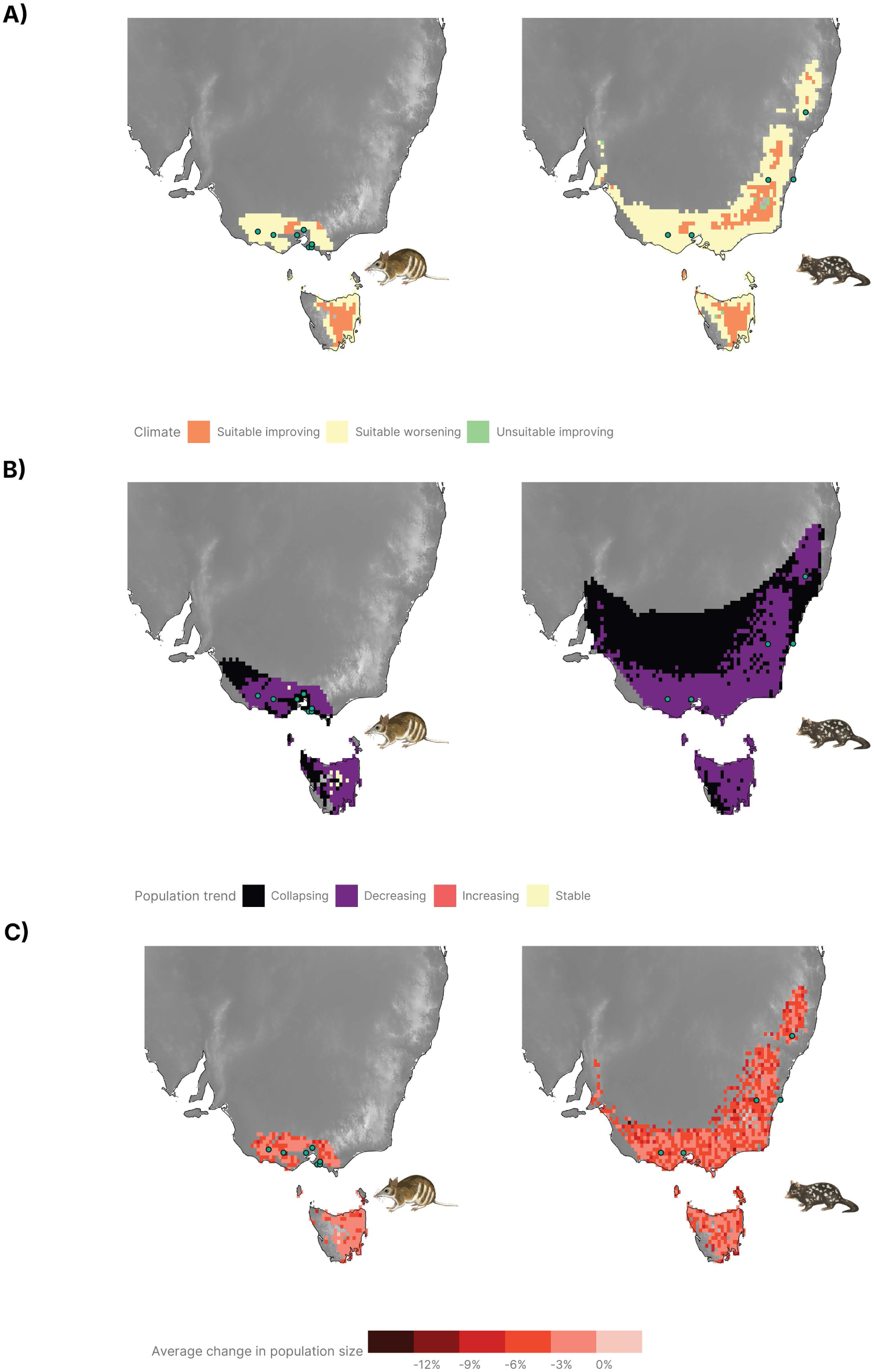
Projections from 2024 to 2100 for *Perameles gunnii* (left) and *Dasyurus viverrinus* (right). A) Trends in climate suitability, based on a linear regression of ensemble climate suitability against time for each cell. “Suitable” refers to cells that are currently above the suitability threshold. “Unsuitable” refers to cells that are currently below the suitability threshold. Cells which are below the suitability threshold in 2100 are omitted. B) Trends in population size, based on a linear regression of predicted population size against time for each cell. “Decreasing” refers to cells with a significantly (p < 0.05) negative regression slope, “Increasing” refers to cells with a significantly (p < 0.05) positive regression slope, and “Stable” refers to cells where the regression slope does not significantly differ from zero (p > 0.05). “Collapsing” refers to cells where the population size drops to zero by 2100. Cells where the population size is currently zero are omitted. C) Average change in population size by 2100, compared to the maximum population size during the century, calculated from cell-specific linear regression coefficients. Only shown for cells where the population is not projected to collapse. Circles in all panels represent existing sanctuary populations.

**Figure 2.**
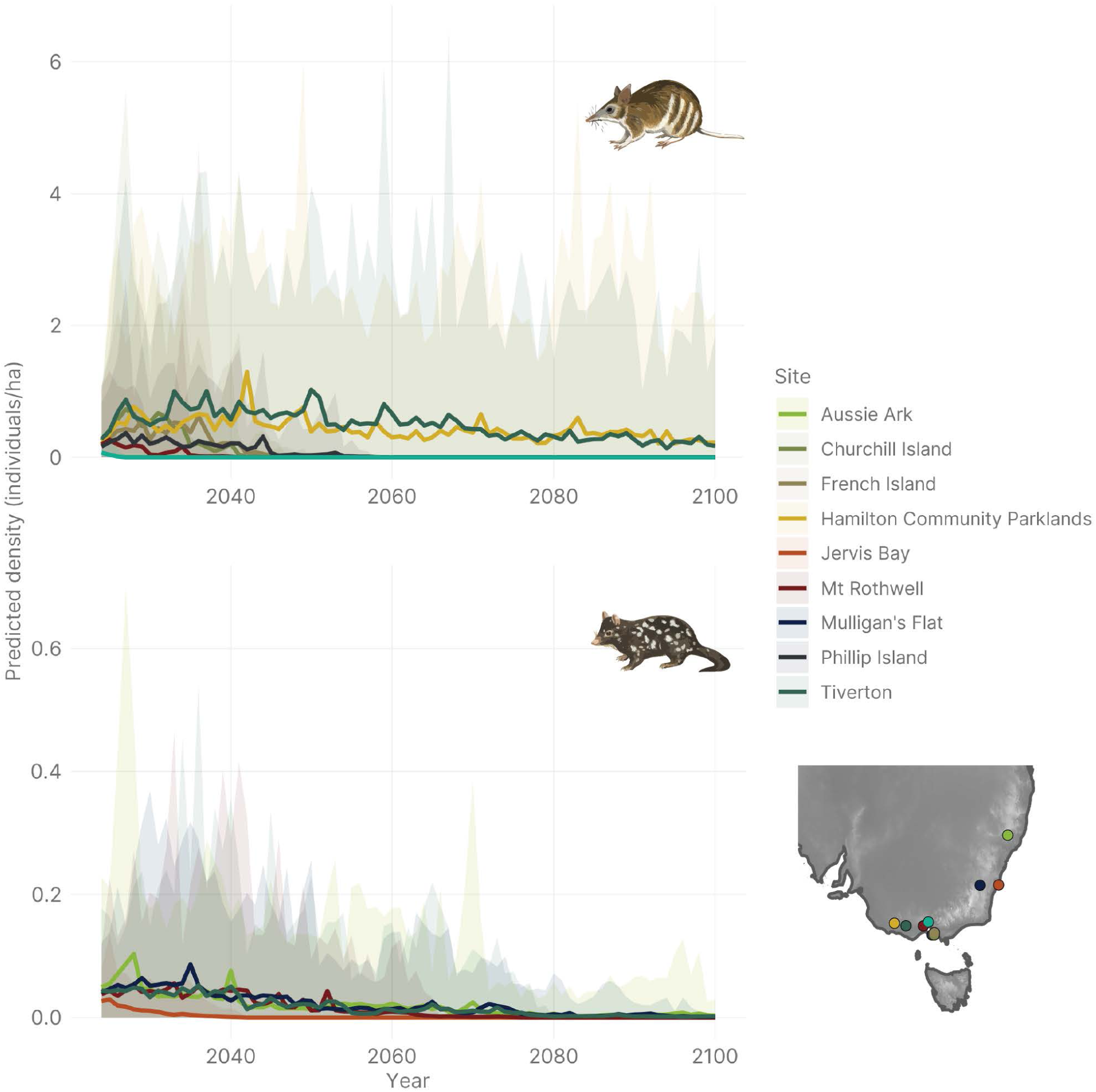
Predicted population trajectories for reintroduced mainland populations of *Perameles gunnii (*top) and *Dasyurus viverrinus* (bottom) in predator-free sanctuaries. The lines show the mean population densities from all 50 simulations, and the shaded areas show the 95% CI around the mean.

These declines closely tracked projected reductions in climatic suitability, but notably, improvements in climatic suitability do not necessarily mean long-term stability or population growth; our models identified many regions with improving climatic suitability, particularly in the Australian Alps, where isolated safe haven populations are nonetheless predicted to decline (Fig. 1B). Rates of decline varied geographically (Fig. 1C), highlighting opportunities for establishing “stressor” safe havens in areas experiencing the fastest projected declines.

These sites could be strategically leveraged to impose controlled environmental stressors that may drive adaptive responses.

Additional modelling scenarios, including those testing alternative assumptions about niche divergence between mainland and Tasmanian populations, are provided in Supporting Information S1 (Figs. S6–10 & S14–15).

### Genetic simulations

Our SEPM simulations suggest that most current safe havens are likely to fail within decades if climatic niches remain static but climate changes. However, evolutionary change has the potential to shift niche limits and improve population persistence under novel conditions. Our genetic simulations show that adaptive variation can rapidly increase in safe haven populations experiencing selection, with stronger selection resulting in faster fitness gains (Fig. 3). This response is driven by rising frequencies of adaptive alleles (Fig. 3B) and an increase in the number of beneficial mutations per individual (Fig. 3C). In contrast, safe haven populations exposed to weak selection exhibited a rapid decline in adaptive allele frequencies, reflecting the combined effects of genetic drift and limited directional selection.

**Figure 3.**
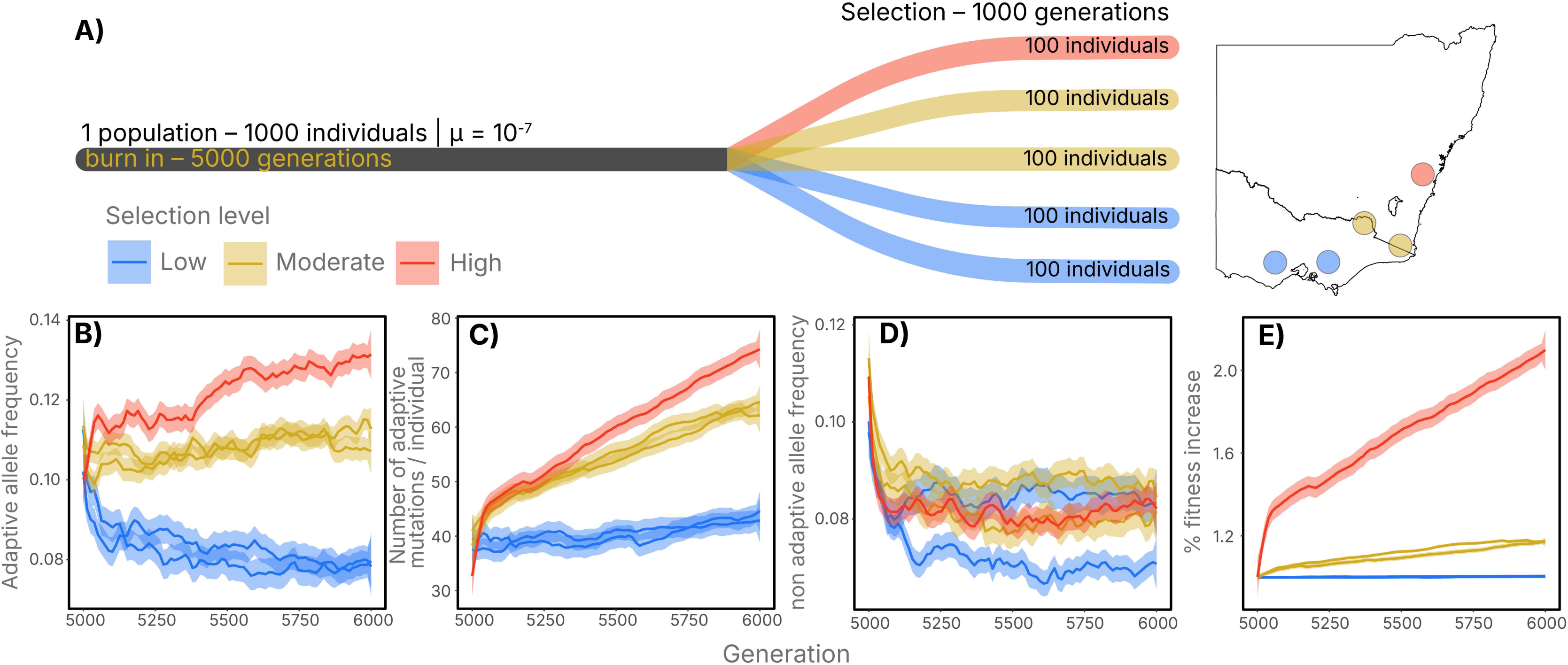
Genetic simulation results. Simulations were sampled every generation over 10 replicate simulations. Panel A shows a schematic of the simulation logic implementing the 500-in-5 framework. Panels B–E show for each selection level (represented by different colours) and generation the mean (bold lines) and 95% CI (shaded area) across all replicate simulations of changes in adaptive (B) and non-adaptive (D) allele frequencies (B), the number of adaptive mutations per individual, and % increase in fitness (E).

These patterns were absent in neutral loci, where allele frequencies stabilised after an initial post-foundation reduction (Fig. 3D), highlighting that the observed increases in fitness were due to selection on adaptive variants rather than random genetic processes. Overall, these results suggest that populations established under stressor conditions, where selection pressures are higher, are likely to accumulate adaptive alleles more rapidly and increase in fitness over time, supporting the rationale of incorporating environmental heterogeneity into the 500-in-5 framework.

### Safe haven predictions

Integrating climate-based population projections with evolutionary outcomes revealed that even modest adaptive responses can influence safe haven population trajectories. When fitness increases from adaptive evolution were incorporated, populations showed moderate to substantial increases in density compared to baseline projections with no adaptive response, although stochastic fluctuations remained evident across all scenarios (Fig. 4). The magnitude of these benefits varied by site and depended on both severity of climatic change and the strength of selection. In some locations, such as Woodlands Historic Park (*P. gunnii*), population declines were too rapid to be offset by fitness gains, whereas in others, including Jervis Bay (*D. viverrinus*), higher fitness increases allowed populations to persist for longer and with higher densities.

**Figure 4.**
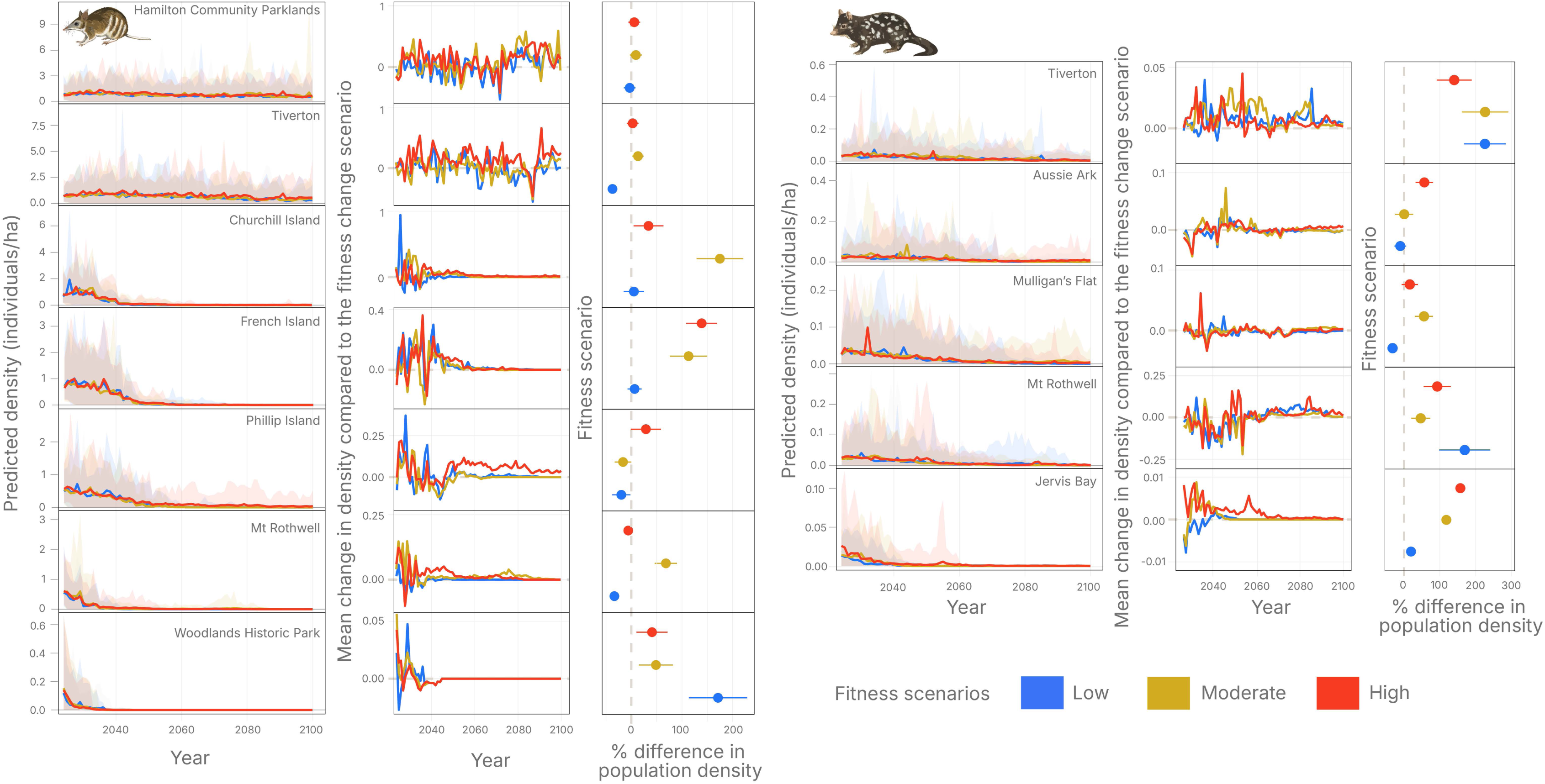
Comparisons of future population trajectories for *Perameles gunnii* (left) and *Dasyurus viverrinus* (right) in current mainland sanctuary populations (see map in Figure 2). Colours represent different fitness scenarios. The leftmost panels for each species show the mean population densities from 50 simulations (bold lines) and the 95% CI (shaded areas). The middle panels show the difference in density in each fitness scenario compared to the baseline simulations with no fitness increases, calculated as the baseline value subtracted from the value in the fitness scenario. The rightmost panels show the mean differences across all timesteps (middle panels), represented as % differences from the baseline values.

Overall, the most pronounced benefits occurred in “stressor” sites, such as Phillip Island and French Island for *P. gunnii* and Mt Rothwell and Tiverton for *D. viverrinus*, where climatic suitability is projected to decline most steeply. In contrast, “core” sites with relatively benign conditions (*e.g.*, Hamilton Community Parklands and Tiverton for *P. gunnii*) showed only minor improvements, reflecting limited selective pressure. These findings demonstrate that incorporating environmental heterogeneity into safe haven design has the potential to enhance adaptive capacity and improve long-term persistence under climate change.

## Discussion

Safe havens are critical tools for preventing species extinctions, often established as insurance populations to safeguard species threatened with imminent extirpation (Hayward *et al*. 2014; Innes *et al*. 2015; Legge *et al*. 2018). Safe havens aim to protect species from ongoing threats, maintain genetic diversity, and provide opportunities for future reintroductions that support long-term persistence within dynamic landscapes (Ringma *et al*. 2018). However, these populations are typically created in isolation, without consideration of metapopulation functionality. As a result, most safe havens focus on short-term survival rather than long-term potential. This *ad hoc* approach can limit evolutionary resilience and reduce the capacity of populations to cope with environmental change (Hayward & Kerley 2009), or even produce unintended outcomes such as loss of anti-predator traits (Evans *et al*. 2022), further limiting the success of future reintroductions. Without integrating evolutionary processes, many safe haven populations are likely to decline or collapse as environmental conditions shift (*e.g.*, Fig. 2). Yet, we demonstrate that deliberately establishing safe havens across heterogeneous environments and managing them as an integrated network can increase adaptive genetic diversity, improve fitness, and enhance population viability.

### Harnessing evolutionary processes for resilience

Polygenic adaptation (small beneficial allele shifts at many loci) can produce rapid fitness gains under directional selection (Barghi *et al*. 2020). When environments change or new selective pressures emerge, populations can shift their mean trait values toward new optima through these subtle, genome-wide allele frequency changes. This allows rapid adaptation, sometimes within just a few generations, provided sufficient SGV is present (Pritchard & Di Rienzo 2010). Polygenic adaptation is particularly effective for complex, quantitative traits because small contributions from many loci can cumulatively generate substantial phenotypic and fitness gains under appropriate selection (Hayward & Sella 2022). Our genetic simulations show that exposing safe haven populations to strong, predictable selective pressures (*e.g.*, climatic stress) can produce measurable fitness increases within tens of generations, provided effective population size remains relatively high (>100 individuals). This supports the potential of the “Goldilocks Zone” approach (enough selection to avoid maladaptation but not too much to cause extinction) to drive adaptation of threatened populations within safe havens (Evans *et al*. 2021, 2022).

Standing genetic variation is a critical determinant of a species’ capacity to persist under changing conditions because it contains alleles that were previously neutral or under weak selection. When environments shift, these alleles may become advantageous—a phenomenon known as conditional neutrality (Mee & Yeaman 2019). Conditionally neutral alleles provide the raw material for rapid adaptation, enabling populations to respond to environmental heterogeneity and shifting selective pressures (Hermisson & Pennings 2005). In a safe haven context, establishing multiple populations with a shared SGV profile but exposed to different selection pressures can accelerate the accumulation of adaptive alleles suited to future conditions.

However, strong selection in stressed safe haven populations can also erode genetic diversity through bottlenecks, drift processes and inbreeding, reducing evolutionary potential if gene flow is not maintained (Lacy 1987). Similarly, populations in benign conditions are prone to losing adaptive genetic variation over time—particularly in the first few generations—when strong selective pressures for adaptive traits are relaxed and otherwise alleles (under stressful or variable conditions) may drift to low frequencies or be lost entirely through genetic drift and founder effects (Guzella *et al*. 2018; Willi *et al*. 2022). Assisted migration could help counteract this erosion (Hogg *et al*. 2024; Shaw *et al*. 2025), preserving adaptive variation in benign “core” populations where selection is relaxed and in stressor populations where diversity may be lost (Guzella *et al*. 2018; Willi *et al*. 2022).

### The 500-in-5 framework in practice

The 500-in-5 framework addresses two core evolutionary principles; (i) *Population size*: maintaining >500 breeding individuals per species preserves long-term evolutionary potential and minimises genetic drift (Franklin 1980; Lande 1995; Franklin & Frankham 1998; Willi *et al*. 2006; Weeks *et al*. 2011; Frankham *et al*. 2014), and (ii) *Environmental heterogeneity*: distributing populations across 5 geographically distinct safe havens facilities local adaption and provides insurance against catastrophic events.

Our modelling illustrates the potential of this approach. Selection on SGV increases population fitness (Fig. 3), which translates into improved population trajectories (Fig. 4). Actively managing connectivity between core and stressor sanctuaries could mimic natural metapopulation dynamics, ensuring gene flow to balance demography (in stressor sites) and facilitate the exchange of adaptive alleles (in core sites). This managed evolving metapopulation could increase resilience and long-term viability, potentially leading to increased future reintroduction success.

There are several practical biological issues that must be considered under our 500-in-5 framework. Larger bodied species with larger home ranges and/or lower population densities (*e.g.*, *D. viverrinus*) will require larger safe havens to reach evolutionary thresholds than smaller species with higher densities (*e.g.*, *P. gunnii*). Additionally, species with shorter generation times will respond more rapidly to selection than long-lived species (Welch *et al*. 2008). Finally, the choice and intensity of stressors must be well understood. While we have chosen climate change to demonstrate the 500-in-5 framework, other stressors could also be considered, but ambiguous or poorly characterised stressors (*e.g.*, variable predator pressure) may limit the effectiveness of selection-based strategies.

## Conclusion

The ultimate goal of safe havens is the reintroduction of self-sustaining populations into wild, unmanaged landscapes. Achieving this requires populations capable of coping with shifting biotic and abiotic challenges. The 500-in-5 framework provides a pathway to achieve this by embedding evolutionary principles into safe haven design and management. Our findings indicate that evolutionary processes must be actively managed; otherwise, populations risk decline even within predator-free safe havens. By strategically distributing populations, fostering gene flow, and leveraging polygenic adaptation, managers can enhance evolutionary potential, improve fitness trajectories, and increase the likelihood of long-term persistence.

Safe haven networks designed in this way are not only insurance against extinction but also incubators of adaptation, ensuring that populations retain the capacity to thrive in dynamic and unpredictable landscapes.

## Acknowledgements

We acknowledge the Traditional Custodians of the land on which our research was conducted and pay our respects to their Elders past and present. We thank Duncan Sutherland for sharing data from long-term eastern barred bandicoot monitoring on Churchill Island. We also thank Annette Rypalski, Dale Crisp and the Odonata team for discussions around eastern barred bandicoots and eastern quolls. This work was supported by the Coexistence Conservation Laboratory (www.coexistenceconservationlab.org) at the Fenner School of Environment and Society at the Australian National University, and a grant from the Amazon Right Now Climate Fund.

The authors declare no conflict of interests.

## Supporting Information S1: Projecting population trajectories to inform selection of sanctuary populations for *Perameles gunnii* and *Dasyurus viverrinus*

### Background

Modelling offers a promising avenue to design sanctuary networks under the 500 in 5 framework and has long been a staple of research in climate change impacts on biodiversity (Pacifici *et al*. 2015). Mechanistic and correlative ecological niche models, also known as habitat suitability models, have been widely used to predict distributional shifts of species under climate change (Pearson & Dawson 2003; Araújo *et al*. 2005). Correlative niche models predict future habitat suitability under novel conditions by estimating a species’ realised climatic niche from historical and contemporary records (Wiens *et al*. 2009). They can be coupled with process-based population models to derive more realistic estimates of population persistence under changing climates, while explicitly incorporating relevant processes such as growth dynamics, dispersal and population connectivity (Keith *et al*. 2008; Fordham *et al*. 2013a).

A recent study (Tomlinson *et al*. 2023) used such a coupled approach to reconstruct the extinction patterns of numbats (*Myrmecobius fasciatus*). Tomlinson *et al*. leveraged historical records to estimate the climatic niche of numbats and infer realistic parameters of population dynamics by using flat priors to run thousands of simulations of interconnected populations across Australia, coupled with modelled predation by introduced cats. Using external validation targets, Tomlinson *et al*. were able to select parameter values which resulted in simulations most closely matching the observed temporal and spatial pattern of the species’ decline. These parameters were then used to run ’counterfactual’ simulations to see how numbat populations might have fared without the introduced predator.

We adapted the approach used by Tomlinson *et al*. (2023) to identify potential sites for a 500 in 5 conservation program for two threatened Australian marsupials, aimed to adapt them to changing climates. Eastern barred bandicoots (*Perameles gunnii*) and eastern quolls (*Dasyurus viverrinus*) both used to be widespread in southeastern Australia. They experienced sharp declines and complete extirpation from the mainland during the 20^th^ century, predominantly due to predation by introduced foxes (*Vulpes vulpes*) (Jones & Rose 2001; Kinnear *et al*. 2002; DSE Victoria 2009), but remnant populations remain in the largely fox-free Tasmania.

Reintroduced populations of *P. gunnii* on the mainland are derived from a captive-bred colony originally sourced from the last remnant mainland population in Hamilton, Victoria (Weeks *et al*. 2013). In part due to the small founder populations (Myorniuk 1993), these reintroduced populations have experienced rapid declines in genetic diversity over the last decades (Weeks *et al*. 2013). Conversely, the original mainland populations of *D. viverrinus* are completely extinct: they disappeared much earlier than *P. gunnii* (1960s vs 1990s) and reintroduced mainland populations are derived solely from translocated Tasmanian animals (Woinarski *et al*. 2014; Wilson *et al*. 2020, 2023). Little is known about the genetics of reintroduced *D. viverrinus* populations. However, there is significant population structure among Tasmanian populations of the species (Firestone *et al*. 2000; Cardoso *et al*. 2014), with reduced genetic diversity in coastal, peripheral populations (Cardoso *et al*. 2014).

Reintroduced populations in predator-free sanctuaries have been established throughout Victoria and New South Wales, with plans to expand the programs to additional sites in the future. These two species are therefore excellent case studies to explore the potential for using managed populations to support evolutionary resilience to climate change. We use the coupled modelling approach to reconstruct the extinction patterns of *P. gunii* and *D. viverrinus* from the Australian mainland and infer realistic parameters of population growth. We then use these parameters in a simulation framework with future projections of climate suitability, but without predation or dispersal, to extrapolate locations across the species’ historical ranges in which predator-free sanctuaries could house populations which would be exposed to a sustainable rate of climate change that will offer an opportunity for adaptation.

### Supplementary Methods

#### Data collection

##### Occurrence data

We downloaded occurrence records for *D. viverrinus* and *P. gunnii* from the Atlas of Living Australia (Belbin *et al*. 2021) using the *galah* package v1.5.0 (Westgate *et al*. 2024) in R v4.2.1 (R Core Team 2022). We used the ALA General profile filter to omit observations with spatial, taxonomic or temporal issues, while maintaining all non-fossil observations post 1700. This allowed us to include both historical (< 1960) and contemporary observations of the target species (Fig. S1), allowing us to get a more accurate approximation of their climatic niches prior to European colonisation and subsequent range contractions.

**Figure S1.**
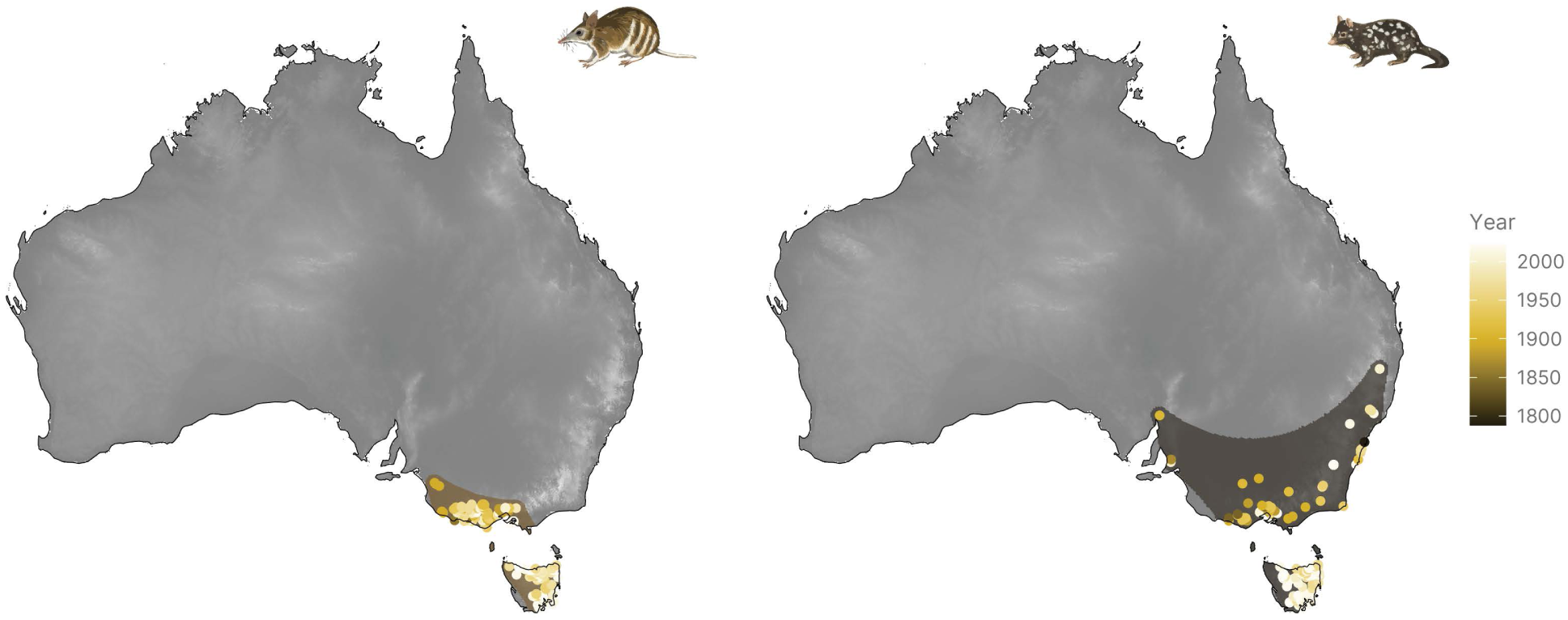
Occurrence records for *Dasyurus viverrinus* ( right) and *Perameles gunnii* ( left) coloured by year of observation. The shaded polygons around the observations in each panel represent the study area for simulations, generated by calculating a 50 km buffer around an α -convex hull calculated around the observations.

In order to reduce computation time for simulations (see below) we minimised the study area from Australia-wide to a large buffer zone around the observations for each species. We did this by calculating an α-convex hull around each species’ observations using the ‘ahull’ function from the *alphahull* package v2.5 (Pateiro-Lopez & Rodriguez-Casal 2022) with an α value of 10 and then applying a buffer of 50 km (Fig. S1).

##### Climate data

We used three climatic variables to construct climatic niches for our target species: average monthly temperatures (°C) for the warmest three months (January, February and March; WQT), average monthly temperatures (°C) for the coolest three months (June, July and August; CQT), and total annual rainfall (mm; PREC). These represent climatic extremes which are relevant to marsupial physiology (Withers *et al*. 2006) and more broadly to physiological and biotic responses to climate change (Bozinovic & Pörtner 2015; Murali *et al*. 2023).

We used annual Australia-wide layers from 1750–2020 generated for these three variables by Tomlinson *et al*. (2023). They used data from PaleoView v1.5.1 (Fordham *et al*. 2017) and the RCP 4.5 projection from the StableClim database (Brown *et al*. 2020) to generate layers at a 0.2° resolution, bias corrected using a delta correction (Beyer *et al*. 2020). We then extracted WQT, CQT and PREC for each observation of *D. viverrinus* and *P. gunnii* based on the coordinates and year of observation, after removing duplicate records.

To allow future projections of climate suitability, we used the same methodology as Tomlinson *et al*. (2023) to extend the RCP 4.5 projections to 2100, capturing predicted climatic changes during the 21^st^ century under a probable baseline “middle-of-the-road” climate change scenario, whereby emissions peak around 2040 then decline, leading to a stabile CO2 concentration in the atmosphere by 2100 (Thomson *et al*. 2011).

#### Climate suitability models

##### Niche estimation and projection

We used four different approaches to estimate the climatic niches of *D. viverrinus* and *P. gunnii*. While the types of models we constructed are often referred to in the literature as habitat suitability models, we refer to them throughout the text as climate suitability models, to reflect that the climatic niche is only one element of the habitat requirements for a species. All four approaches we used are variations of climate envelope models based on presence-only data and so allow us to reconstruct the climatic niches of our target species through time (Nogués-Bravo 2009):

1. Hypervolume niche model: The niche of each species was estimated as a 3-dimensional hypervolume (Blonder *et al*. 2014) using the *hypervolume* package v3.1.1 (Blonder *et al*. 2023). We used Gaussian kernel density estimation (KDE) with the bandwidth estimated using the Silverman estimator (Silverman 1986). Hypervolumes were then projected onto annual climatic layers using the ‘hypervolume_project’ function and we rescaled all layers to range between 0 to 1, using the 95^th^ percentile of climate suitability at any time in the study region as the upper value.
2. Range bagging: The niche of each species was estimated using range bagging (Drake 2015), in essence a method of climatic bootstrapping. Over 100 iterations, two out of three dimensions (climatic predictor variables) were randomly chosen and 50% of observations were sampled. We then calculated the smallest convex hull around each subsample using the ‘convhulln’ function from the *geometry* package v0.4.6.1 (Habel *et al*. 2022). We calculated a total suitability score for each cell in each year as the proportion of convex hulls within which the cells’ climatic conditions lie, calculated using the ‘inhulln’ function from the same package.
3. Weighted range bagging: The niche of each species was estimated in a similar manner to the range bagging algorithm described above. However, in this approach, we used a weighted sampling approach, where the probability of sampling each observation was inversely proportional to the number of observations in that year. This gave a higher probability of sampling historical observations, in order to prevent biased niche estimates due to the fact that contemporary observations are much more common in the dataset.
4. Ensemble: Since suitability scores from all three approaches range from 0 to 1, we calculated the mean suitability score for each cell to generate an ensemble climate suitability score.

We used the four approaches to explore two distinct scenarios. In the first (‘Overlapping niches’), we assumed there is complete overlap in fundamental niches between populations on the mainland and in Tasmania. We therefore used all observations to construct the climate suitability models. In the second (‘Divergent niches’), we assumed that enough time has passed since the separation of Tasmanian from mainland populations for their fundamental niches to diverge completely. We therefore constructed two separate models, one for the ‘Mainland niche’ using only observations from the mainland, and one for the ‘Tasmania niche’ using only observations from Tasmania.

##### Performance and threshold estimation

For all four approaches and two scenarios we estimated model performance by calculating the continuous Boyce index (Hirzel *et al*. 2006), a measure of model performance for presence-only habitat suitability models. The Boyce index varies from -1 to 1; positive values indicate model predictions that are consistent with observed presence records; values close to zero indicate the model is not different from a random model; and negative values indicate an incorrect model which predicts low suitability for areas with frequent presences. We calculated the Boyce index for each model using the projected climate suitability layers and the downloaded occurrence records with the ‘ecospat.boyce’ function from the *ecospat* package v4.0.0 (Habel *et al*. 2022). We used all occurrence records to calculate the index for models in the ‘Overlapping niches’ scenario, but in the ‘Divergent niches’ scenario we only used observations from the mainland to calculate the index for ‘Mainland niche’ models and only observations from Tasmania to calculate the index for ‘Tasmania niche’ models.

We also calculated a suitability threshold for each model as the minimum climate suitability value for which the *predicted-to-expected* (*P*/*E*) ratio *Fi* is higher than 0, *i.e.* the lowest climate suitability value for which there are more occurrence records than predicted by chance. This threshold value was used as a cut-off to designate a cell as suitable or not suitable for the species (see below).

Since the ensemble models generally performed better than the individual models (Table S1), we used the results obtained from the ensemble models in the process-based population models below.

**Table S1.**
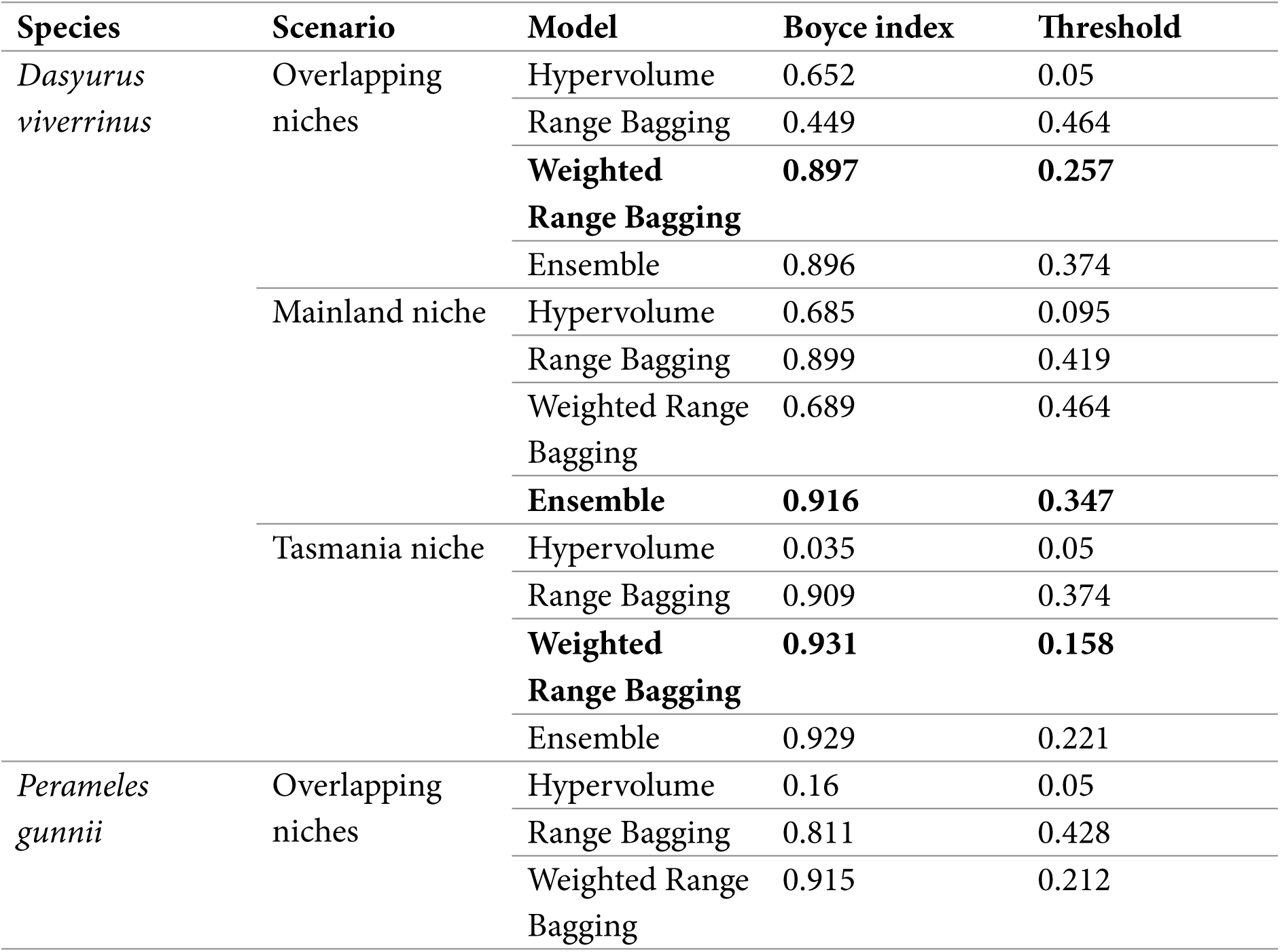
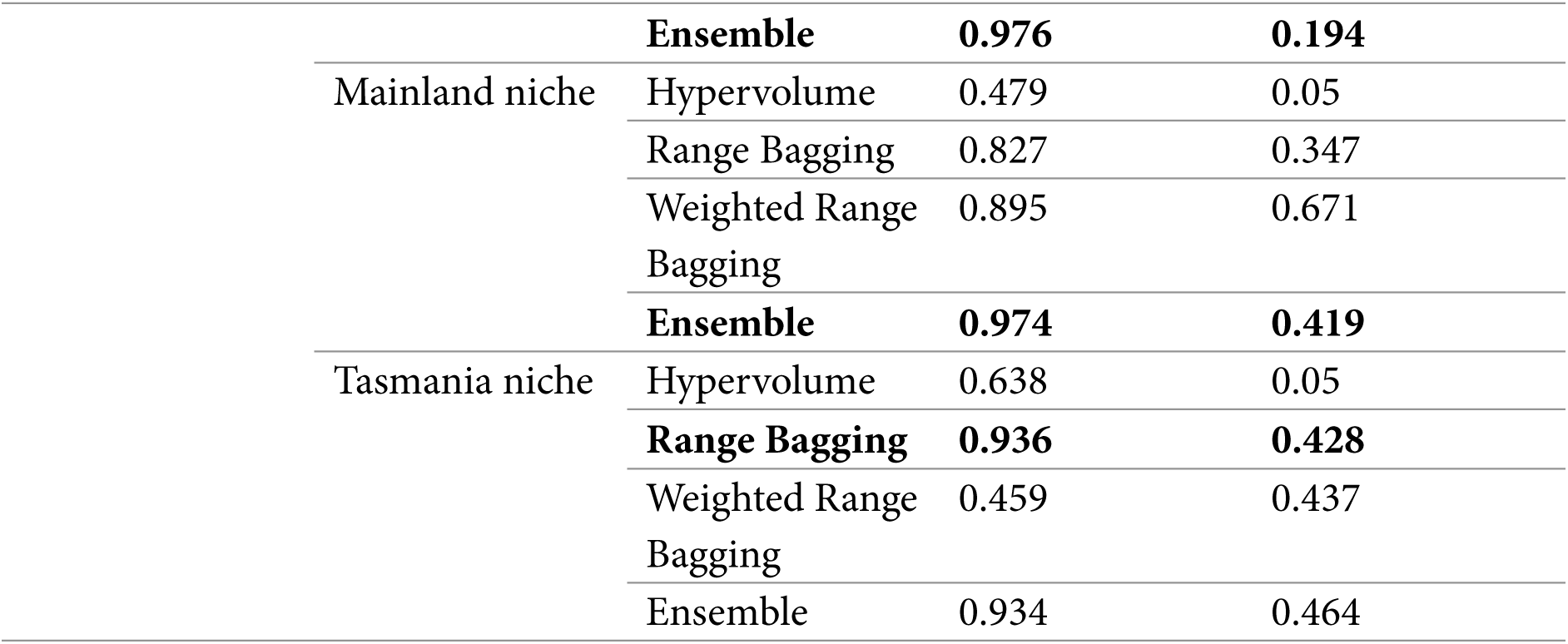
Model performance of climate suitability models. The best performing model (highest value of the Boyce index) for each species and each scenario is in bold. Threshold represents the estimated threshold value for conversion of cells to binary presence/absence.

#### Process-based population models

##### Model structure

We implemented the spatially explicit population modelling (SEPM) framework developed by Tomlinson *et al*. (2023) to simulate landscape-level population processes integrated with climate suitability models. In short, each 20 km x 20 km grid cell in our study area (Fig. S1) is modelled with a single age-stage stochastic population model simulating survival, fecundity, density-dependent population growth, and o@ake (Fordham *et al*. 2021). Dispersal dynamics allow movements of propagules between cells, and carrying capacity of cells is modified based on modelled climate suitability, creating a landscape-wide metapopulation with source-sink dynamics.

##### Population parameter estimates

In order to capture uncertainty around survival, fecundity and population growth of *D. viverrinus* and *P. gunnii*, we generated prior distributions of several parameter estimates for our models based on empirical data (Table S2). In the SEPM, population growth is modelled as a density-dependent process using a Ricker logistic response (Ricker 1954), so that population growth slows as populations approach carrying capacity, reflecting the reduction in available resources per individual. The maximum rate of population increase (*growth_rate_max*) and the maximum carrying capacity in a 20 km x 20 km grid cell (*density_max*) were estimated based on time series data of *D. viverrinus* and *P. gunnii* from sites without foxes (Fig. S2). For both species we estimated these parameters by fitting a Ricker logistic growth model to time series population data using a generalised linear model with a Poisson distribution and a log-link function and setting flat priors around the estimated values (Table S2). For *P. gunnii*, we estimated parameters based on a time series of surveys from a re-introduced population in the predator-free Churchill Island, Victoria (D Sutherland, pers. comm.). For *D. viverrinus*, we estimated parameters based on a time series dataset of surveys of a population in Cradoc, Tasmania, where foxes do not occur (Godsell, 1983).

**Figure S2.**
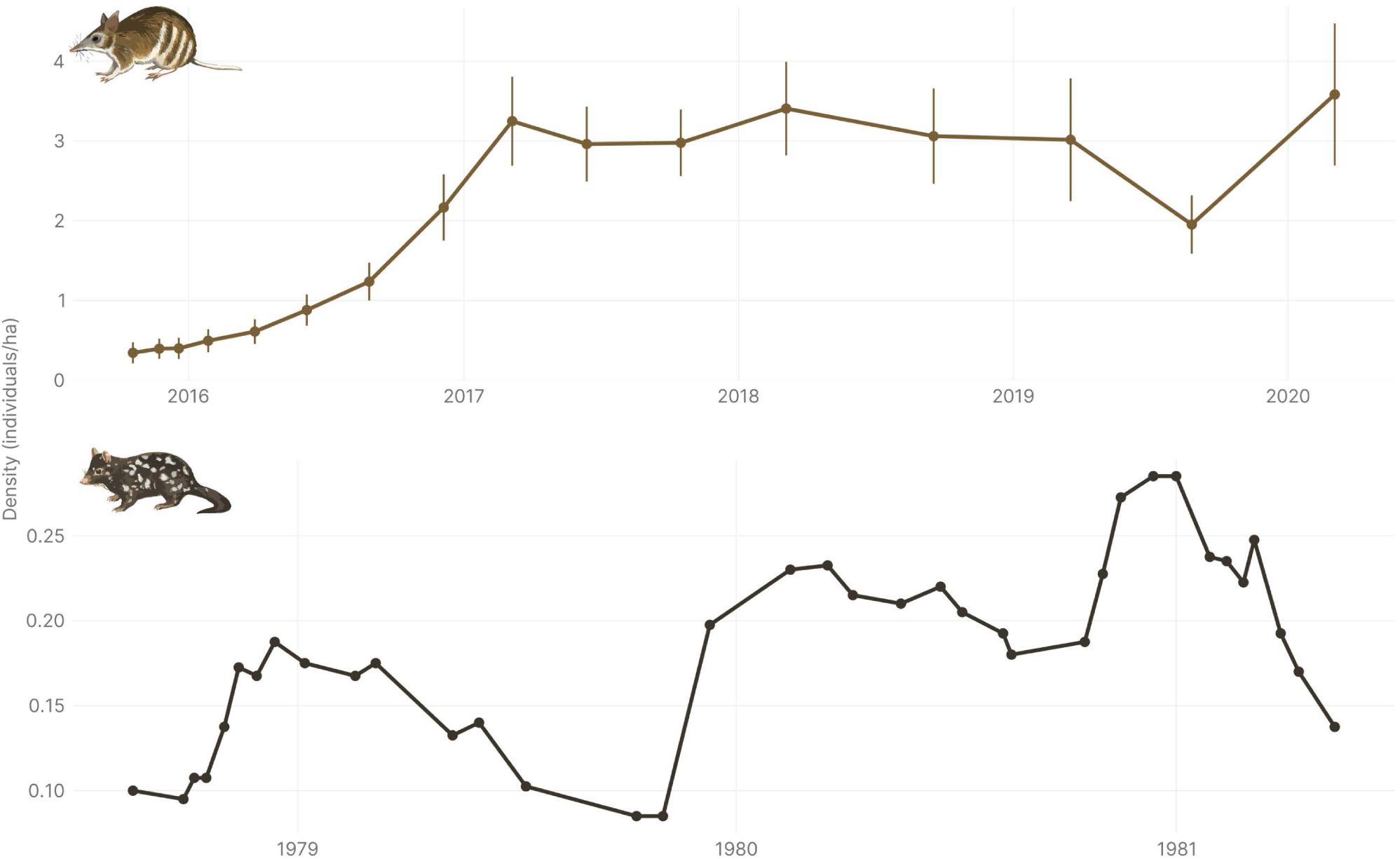
Time series population data of *Perameles gunnii* (top) from a re-introduced population in Churchill Island, Victoria (D Sutherland, pers. comm.) and *Dasyurus viverrinus* (bottom) from a population in Cradoc, Tasmania (adapted from Godsell 1983).

**Table S2.** Parameter values used for process-oriented modelling. Fixed values were consistent across all simulations, whereas variable parameters varied across the prior parameter space sampled via Latin-Hypercube sampling.

| Parameter | type | *Dasyurus viverrinus* | *Perameles gunnii* |
| --- | --- | --- | --- |
| **Demographic parameters** |  |  |  |
| *growth_rate_max* | Variable | 0.05–0.73 | 0.10–1.47 |
| *standard_deviation* | Variable | 0.00–1.46 | 0.00–1.47 |
| *density_max* | Variable | 0.082–0.163 / ha (3,262–65,214 individuals per grid cell) | 1.14–2.29 / ha (45,767–91,534 individuals per grid cell) |
| **Dispersal parameters** |  |  |  |
| *dispersal_max* | Fixed | 20 km | 20 km |
| *dispersal_p* | Variable | 0.001–0.600 | 0.001–0.600 |
| *dispersal_b* | Variable | 0–20 km | 0–6 km |
| **Allee effect** |  |  |  |
| *abundance_threshold* | Variable | 0.004–0.008 / ha (163–326 individuals per grid cell) | 0.057–0.114 / ha (2,288–4,577 individuals per grid cell) |
| *occupancy_threshold* | Fixed | 10 cells | 10 cells |
| **Predation parameters** |  |  |  |
| *harvest_max* | Variable | 0.001–0.850 | 0.001–0.850 |
| *harvest_z* | Variable | 1–2 | 1–2 |

**Table S3.** Mainland sites with reintroduced populations of *Dasyurus viverrinus* and *Perameles gunnii* in predator-free sanctuaries.

| Site | Latitude | Longitude | State | *Dasyurus viverrinus* | *Perameles gunnii* |
| --- | --- | --- | --- | --- | --- |
| Mt Rothwell | -37.8966 | 144.4418 | Victoria | Present | Present |
| **Tiverton** | -37.8879 | 142.9832 | Victoria | Present | Present |
| **Mulligan's Flat** | -35.1656 | 149.1628 | Australian Capital Territory | Present | Absent |
| **Jervis Bay** | -35.1429 | 150.7121 | New South Wales | Present | Absent |
| **Aussie Ark** | -31.8219 | 151.4666 | New South Wales | Present | Absent |
| **Phillip Island** | -38.4994 | 145.1627 | Victoria | Absent | Present |
| **Churchill Island** | -38.4999 | 145.3397 | Victoria | Absent | Present |
| **French Island** | -38.3514 | 145.3489 | Victoria | Absent | Present |
| **Woodlands Historic Park** | -37.6332 | 144.8512 | Victoria | Absent | Present |
| **Hamilton Community Parklands** | -37.7208 | 142.0262 | Victoria | Absent | Present |

Carrying capacity of cells was based on modelled climate suitability (VanDerWal *et al*. 2009). The values of *density_max* for both species were multiplied by 0.75 (Tomlinson *et al*. 2023) based on the assumption that not the entirety of the cell will be suitable climate (Fordham *et al*. 2013b). *density_max* values for each cell in each year were then multiplied by its climate suitability score.

For each cell we set a value of *abundance_threshold* to represent a local Allee effect: “a positive relationship between any component of individual fitness and either numbers or density of conspecifics” (Stephens & Sutherland 1999; Stephens *et al*. 1999). If at any time in the simulation the number of individuals in the cell reached *abundance_threshold* the cell abundance would be made zero, representing local extirpation of the population (Fordham *et al*. 2013b). For both species these values were set as 5% of the maximum carrying capacity (Table S2). We also fixed *occupancy_threshold*, the number of occupied cells below which simulations were terminated (total extirpation), at 10, representing a “global” Allee effect.

Since we lacked reliable data on dispersal capability of *D. viverrinus* and *P. gunnii*, we used the same parameters Tomlinson *et al*. (2023) set for their numbat model, where dispersal rates are modelled using a distance-based function based on an average dispersal distance (*dispersal_b*), which we set to vary between 0 and 20 km for *D. viverrinus* (Wilson *et al*. 2020) and 0 and 6 km for *P. gunnii* (Dufty 1991). We fixed the maximum dispersal distance (*dispersal_max*) at 20 km, the resolution of cells in the grid.

##### Fox predation

Predation by foxes is considered one of the main causes for the extirpation of *D. viverrinus* and *P. gunnii* from mainland Australia (Johnson 2006; Woinarski *et al*. 2019). We therefore modelled predation by foxes as a harvest term in our SEPM, similar to how Tomlinson *et al*. (2023) treated cat predation. We used Australia-wide layers of fox abundance estimated from surveys of fox density (Stobo-Wilson *et al*. 2022). We also used similar values of *harvest_z*, a value representing how far predation is from the maximum rate, and *harvest_max*, a value representing the maximum harvest rate (proportion of the prey population consumed by predators).

Finally, harvest rate per cell is also a function of timing of fox arrival. We therefore generated a raster layer of fox arrival timing per cell in Australia by linear interpolation from contours based on a published map of fox spread in Australia (Fairfax, 2019; Fig. S3).

**Figure S3.**
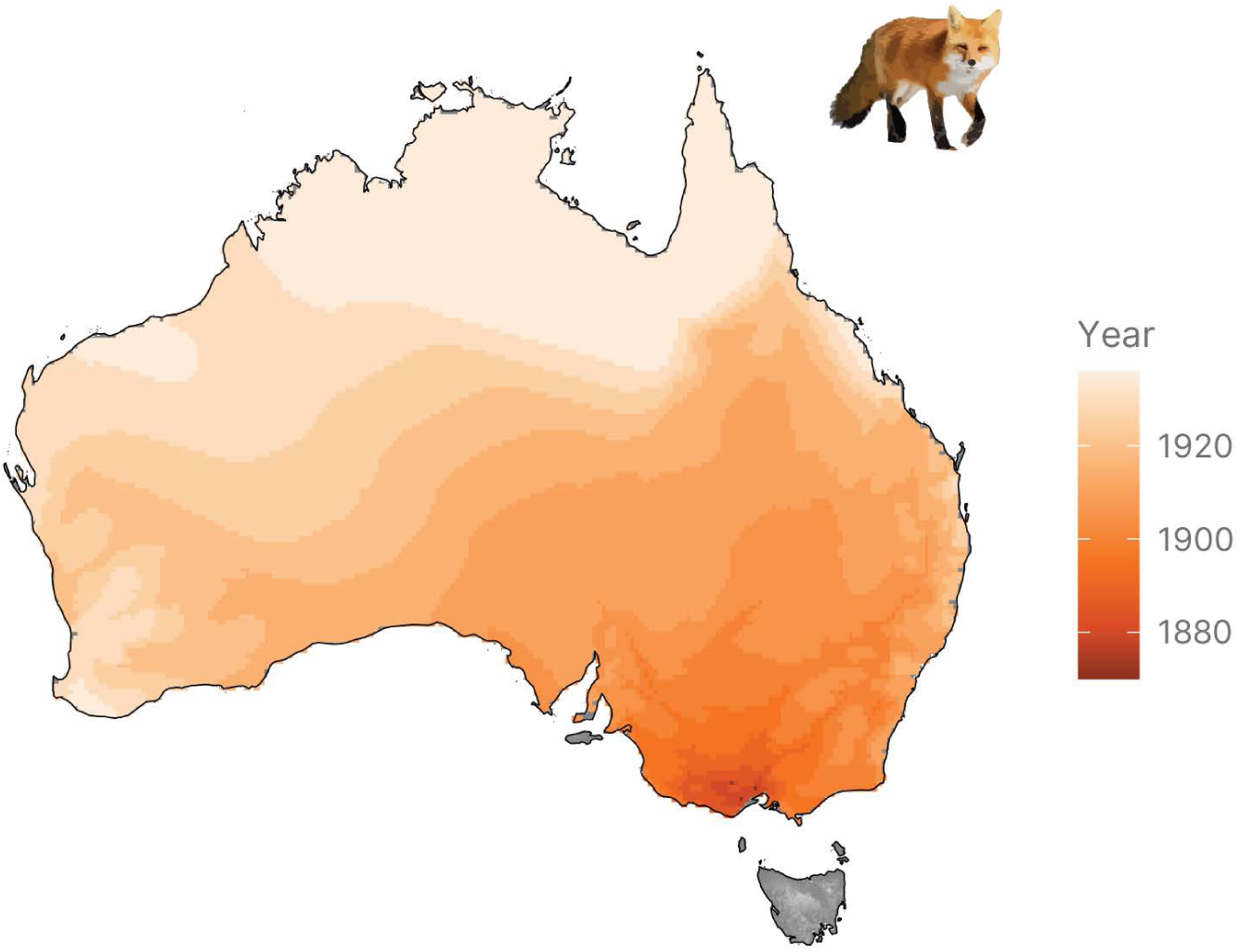
Map of timing of red fox (*Vulpes vulpes*) spread across Australia, adapted from Fairfax (2019) to a 20 x 20 km grid. Note that foxes have not spread to Tasmania.

#### Simulations of future population trajectories

##### Parameter estimation

We used a pattern-oriented modelling approach with the *paleopop* package v2.1.5 (Haythorne *et al*. 2024) to simulate scenarios to match empirically derived validation targets (van der Vaart *et al*. 2015). We generated flat prior distributions of parameter estimates based on empirical data on *D. viverrinus* and *P. gunnii* population dynamics (Table S2) using Latin-Hypercube sampling (Prowse *et al*. 2016) to create 20,000 models with different combinations of parameter values.

We then ran all 20,000 simulations at annual time steps for 351 years (1750–2100) following a climatically stable burn-in period of 50 years to allow initial population abundances to stabilise prior to the introduction of climatic changes and fox predation.

We then used the ‘abc’ function from the *abc* package v2.2.1 (Csilléry *et al*. 2012) to perform parameter estimation using approximate Bayesian computation. Based on the extirpation pattern of both species across the Australian mainland (*D. viverrinus* extirpated from South Australia by 1910, from Victoria by 1950, and from New South Wales by 1965; *P. gunnii* extirpated from South Australia by 1900, and from Victoria by 1990) we calculated the following validation targets: the proportion of known observation records populated by the model, the proportion of extirpation targets met (score of 1 for each mainland population extirpated, divided by the number of populations: 3 for *D. viverrinus* and 2 for *P. gunnii*), a timing penalty for extirpation (calculated as the mean of the absolute difference in years between the time of extirpation from each state in the simulation and the historical time of extirpation), and the status of the Tasmanian population at 2023 (1 if extant, 0 if extirpated). The independent validation targets were defined as 1 (proportion of known observation records populated), 1 (proportion of extirpation targets), 0 (timing penalty), and 1 (Tasmanian population status). We calculated the target metrics for each simulation and used the ‘rejection’ method to retain the 50 simulations (0.25%) which were closest to the independent validation targets based on Euclidean distance in the target metrics.

##### Future simulations

We used the estimated parameter values (Fig. S4) to run 50 new simulations running at annual time steps for 77 years (2024–2100), simulating for each cell how a population of the target species would fare under climate change in a predator-free sanctuary. Therefore, we divided *abundance_threshold* and *density_max* by 30 and by 0.75 to simulate a 1500 ha sanctuary of suitable habitat, set *occupancy_threshold* to 0, and disabled dispersal (by setting *dispersal_max* to 0 km) and fox predation (by setting *harvest = FALSE* in the ‘PaleoPopModel’ object). Thus, we used realistic parameter estimates for survival, fecundity and population growth of our target species, derived from simulations which recreated fox-mediated extirpation from the mainland, to simulate how climate change is going to affect the survival potential of the species in closed populations without the presence of introduced predators.

**Figure S4.**
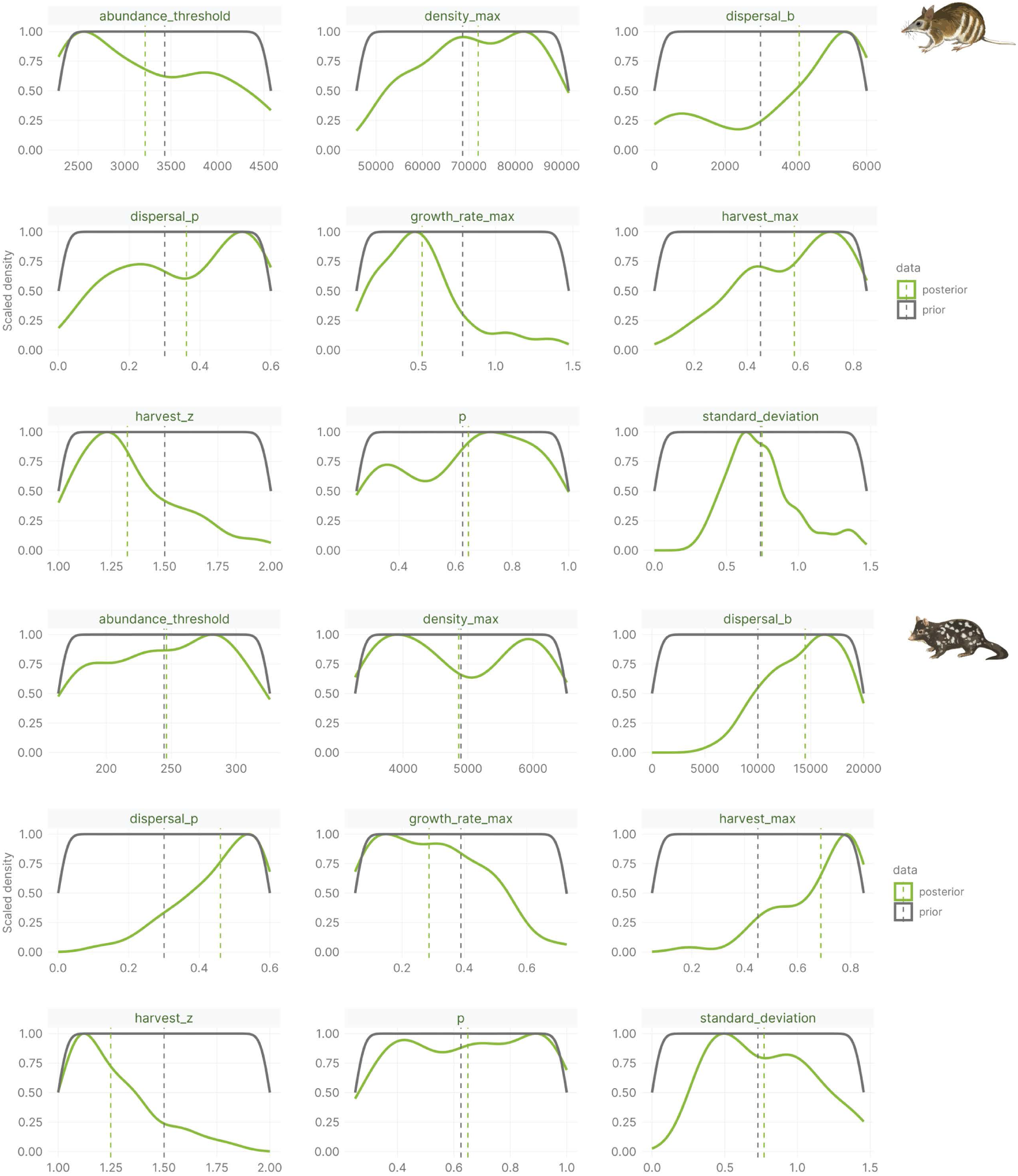
Parameter values as estimated by approximate Bayesian computation for *Perameles gunnii* (left) and *Dasyurus viverrinus* (right). The grey lines represent the prior distributions of parameter values used in 20,000 simulations and sampled using Latin-Hypercube sampling. The green lines represent the posterior distributions of parameter estimates from the 50 simulations closes to the validation targets.

We used initial population sizes of 100 animals for *P. gunnii* and 25 animals for *D. viverrinus*, to reflect the overall lower population densities of the latter. These simulations were run under both scenarios (‘Overlapping niches’ and ‘Divergent niches’) to explore how different assumptions about niche evolution affect projected population trajectories.

Using the simulations across the entire study area, for each cell with a population size above the abundance threshold in 2023 we ran a linear regression of relative population size (calculated as a proportion compared to the maximum population size at any time step) against time (ranging from 2024 to 2100). We then assigned each cell a value of ‘Stable’ (regression slope not significantly different from 0; *p* > 0.05), ‘Increasing’ (regression slope significantly positive; *β* > 0, *p* < 0.05), ‘Decreasing’ (regression slope significantly negative; *β* < 0, *p* < 0.05), or ‘Collapsing’ (population size reaches 0 by 2100). We also extracted the regression slope for each cell, which represented the average annual change in population size. Multiplying this value by the length of the simulation (77 years) gives us the average overall change in population size by 2100.

We ran one other set of simulations, using the climate suitability values only from the ‘Overlapping niches’ scenario. In this set, we explored three different scenarios of change in climatic suitability, reflecting a fitness increase (low, moderate and high) due to adaptation to changing conditions—the intended outcome by implementing a 500-in-5 program across a climatic gradient in the landscape. For each cell and year, we increased its simulated climatic suitability by 0.001 (low fitness increase scenario), 0.002 (moderate fitness increase scenario) or 0.003 (high fitness increase scenario) multiplied by the number of years since the start of the simulation, up to a maximum climatic suitability value per cell of 1. These translate to annual increases of fitness of 0.1%, 0.2% and 0.3%, respectively, or a cumulative fitness increase of 7.6%, 15.2% and 22.8% by the end of the century.

We then explored population trajectories for *D. viverrinus* and *P. gunnii* which have been reintroduced in predator-free sanctuary sites since their extirpation from the mainland (Table S3). We sampled the projected population trajectories from our future simulations in each site to estimate projected population trajectories by the end of the century in these existing sanctuaries under all simulated scenarios. We compared the change in fitness scenarios to a baseline, no change in fitness scenario, represented by simulations under the ‘Overlapping niches’ scenario, to explore how population trajectories of sanctuary populations might change if they adapt to novel climatic conditions.

### Supplementary Results

#### Climate suitability

Both *D. viverrinus* and *P. gunnii* are predicted to experience a reduction in climate suitability across much of their ranges by the end of the 21^st^ century, coupled with a decline in suitable climate area (Fig. S5). For *P. gunnii* this will be reflected in a rapid decline in high suitability cells (Fig. S5A), leading to an overall decrease in mean climate suitability across the entire study area (Fig. S5B) as mostly low suitability cells remain. In contrast, *D. viverrinus* is predicted to experience a much sharper decline in available climate area (Fig. S5C), while its mean climate suitability remains relatively stable (Fig. S5B), likely driven by the loss of intermediate suitability cells as the distribution of climate suitability becomes strongly bimodal, with peaks at both high and extremely low suitability (Fig. S5A). The differences in the two species’ responses are also evident in the fact that initially, mean climate suitability is higher for *P. gunnii*, but with its projected sharp decline during the 21^st^ century, it is predicted to fall below that of *D. viverrinus* in the latter half of the century (Fig. S5B).

**Figure S5.**
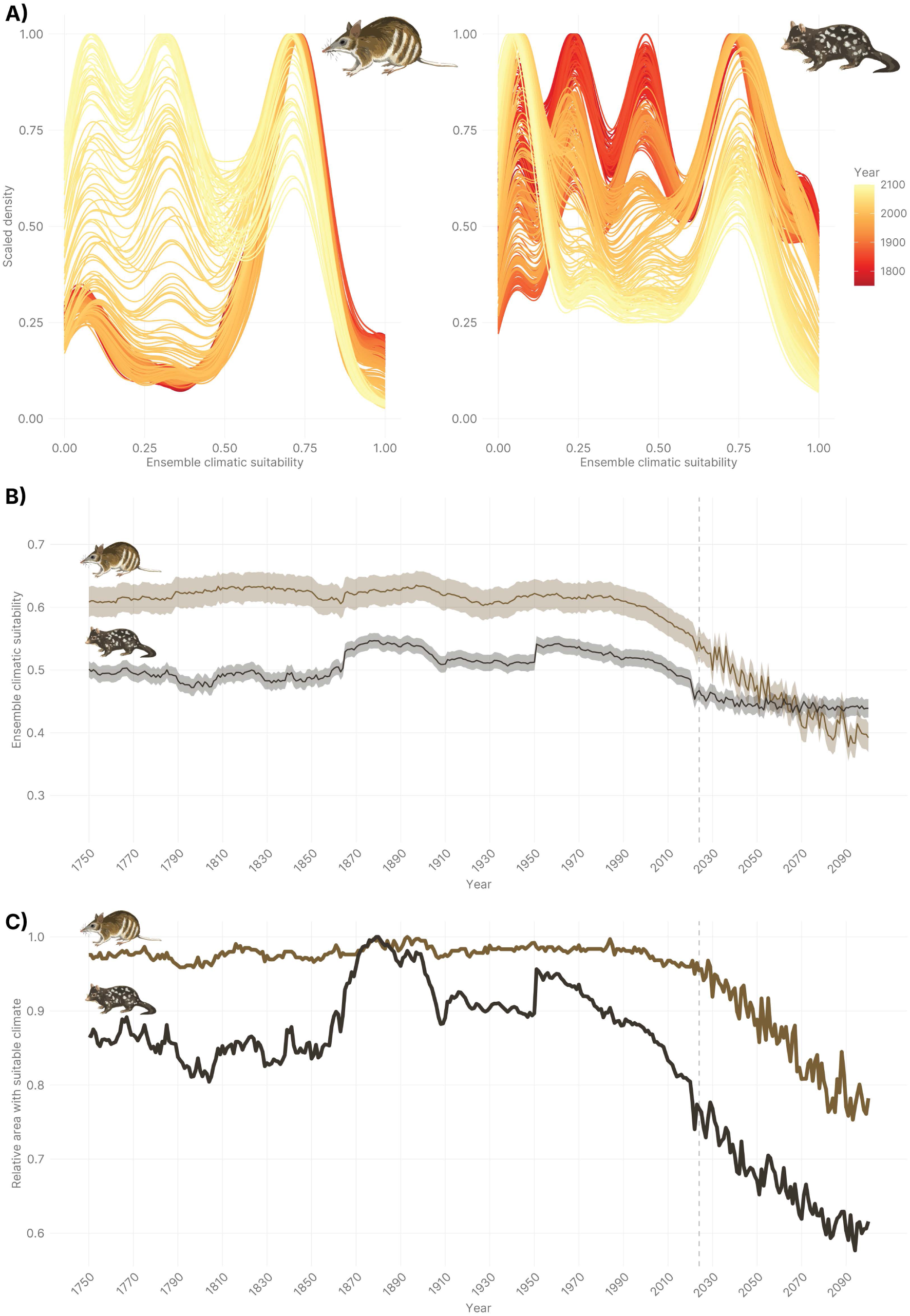
Climate suitability for *Dasyurus viverrinus* and *Perameles gunnii*. A) Distributions of climate suitability scores across the study area from 1750–2100. B) Mean climate suitability across the study area through time, weighted by cell area and omitting cells with a value of 0. Shaded areas represent 95% CI around the mean. C) Sum of suitable climate area through time, defined as all cells with climate suitability above the threshold value. All values are scaled to the maximum area per species across the entire time series. The vertical dashed lines in panels B and C represent the year 2024.

Suitable climate area for *P. gunnii* was historically mostly stable, only beginning to decline towards the end of the 20^th^ century. Meanwhile, suitable climate area for *D. viverrinus* peaked around 1880 followed by a decline, especially since 1950 (Fig. S5C). These declines are not homogenous across the entire study area—for *P. gunnii* climate suitability has been declining steadily in the Victorian grasslands, for *D. viverrinus* climate suitability has been declining steadily in the lowlands and coastal areas, and for both species climate suitability has been increasing in inland Tasmania (Fig. S6). In the latter areas climate suitability is predicted to continue to increase until the end of the century.

**Figure S6.**
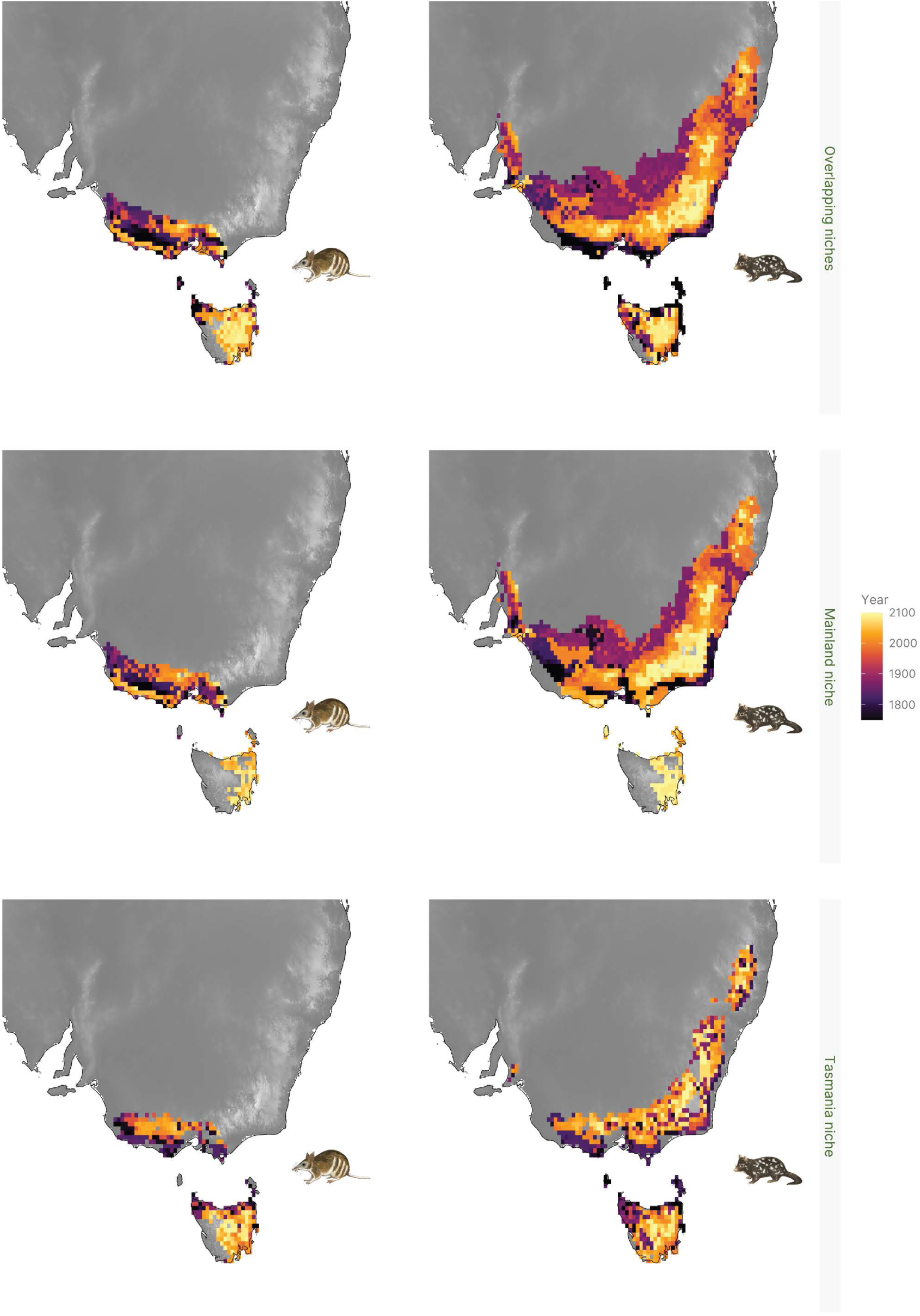
Latest year maximum ensemble climate suitability was reached in each cell for *Perameles gunnii* (left) and *Dasyurus viverrinus* (right) under the two scenarios, ‘Overlapping niches’ and ‘Divergent niches’. Cells where the climate suitability score never exceeded the threshold values are omitted.

These patterns are mostly consistent regardless of the method of climate suitability modelling used or of whether niches are assumed to be completely overlapping or completely divergent (Figs. S7–S10). However, a key difference is that under the ‘Mainland niche’ declines in climate suitability and suitable climate area are predicted to be sharper, whereas under the ‘Tasmania niche’ climate suitability values across the entire study area are predicted to be lower, and in many cases fall below the suitability threshold. This reflects the fact that, despite large overlap in PREC and CQT values, there is almost no overlap between WQT values of Mainland and Tasmanian populations (Fig. S11)—conditions on the mainland are decidedly warmer and somewhat drier than in Tasmania. Under projected climate change, conditions are expected to become warmer throughout Australia, meaning the mainland will become even more climatically extreme whereas Tasmania will become more similar to current conditions on the mainland. Almost all climate suitability models produce qualitatively similar projections, but hypervolume models generally predict larger areas of suitable climate due to these models’ lower threshold values (Table S1).

**Figure S7.**
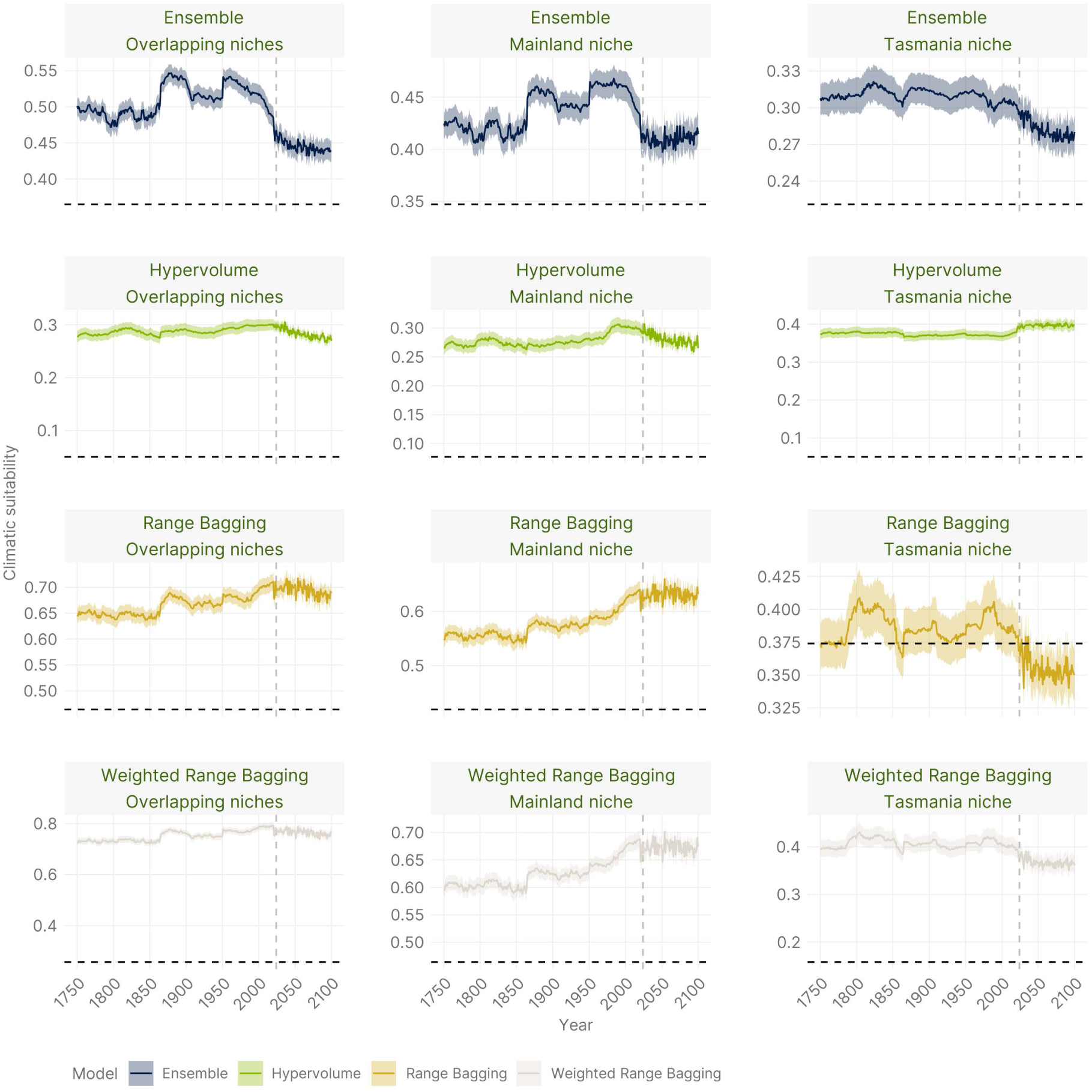
Mean climate suitability across the study area through time for *Dasyurus viverrinus* across all scenarios and climate suitability models, weighted by cell area and omitting cells with a value of 0. Shaded areas represent 95% CI around the mean. The vertical dashed lines represent the year 2024, and the horizontal dashed lines represent the suitability threshold values. Note the y axes have different scales in each panel.

**Figure S8.**
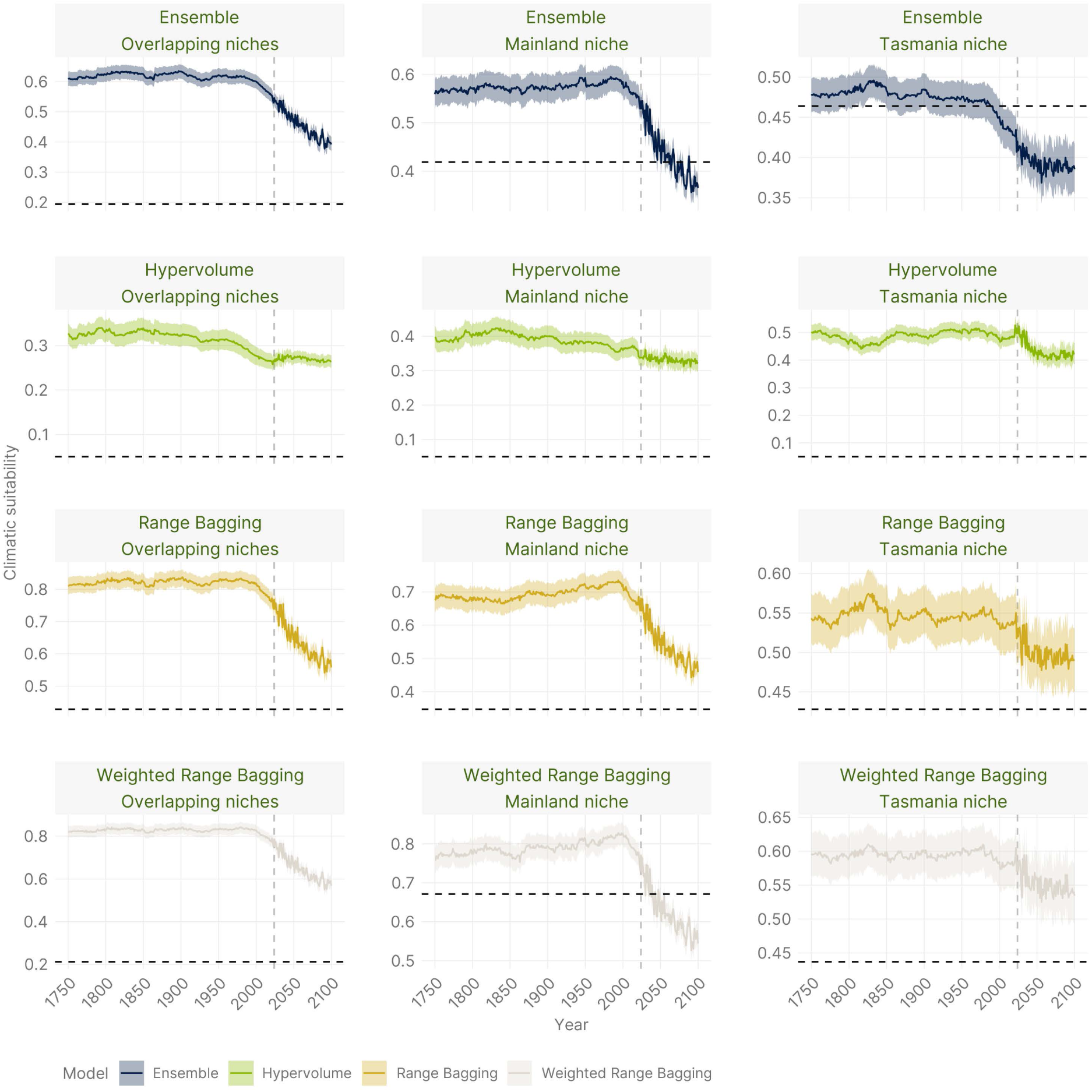
Mean climate suitability across the study area through time for *Perameles gunnii* across all scenarios and climate suitability models, weighted by cell area and omitting cells with a value of 0. Shaded areas represent 95% CI around the mean. The vertical dashed lines represent the year 2024, and the horizontal dashed lines represent the suitability threshold values. Note the y axes have different scales in each panel.

**Figure S9.**
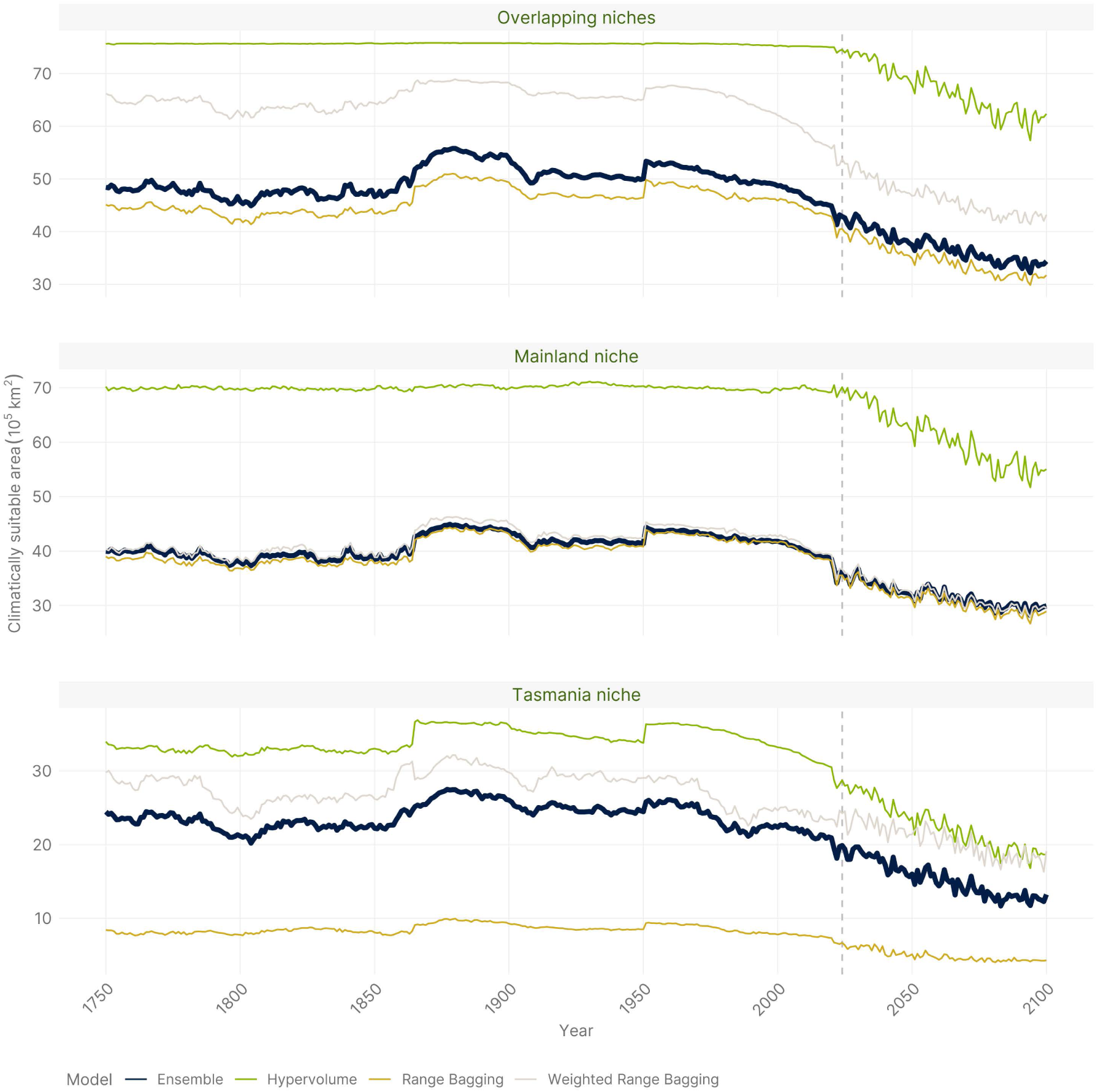
Sum of suitable climate area through time, defined as all cells with climate suitability above the threshold value for *Dasyurus viverrinus* across all scenarios and climate suitability models. The vertical dashed lines represent the year 2024. Note the y axes have different scales in each panel.

**Figure S10.**
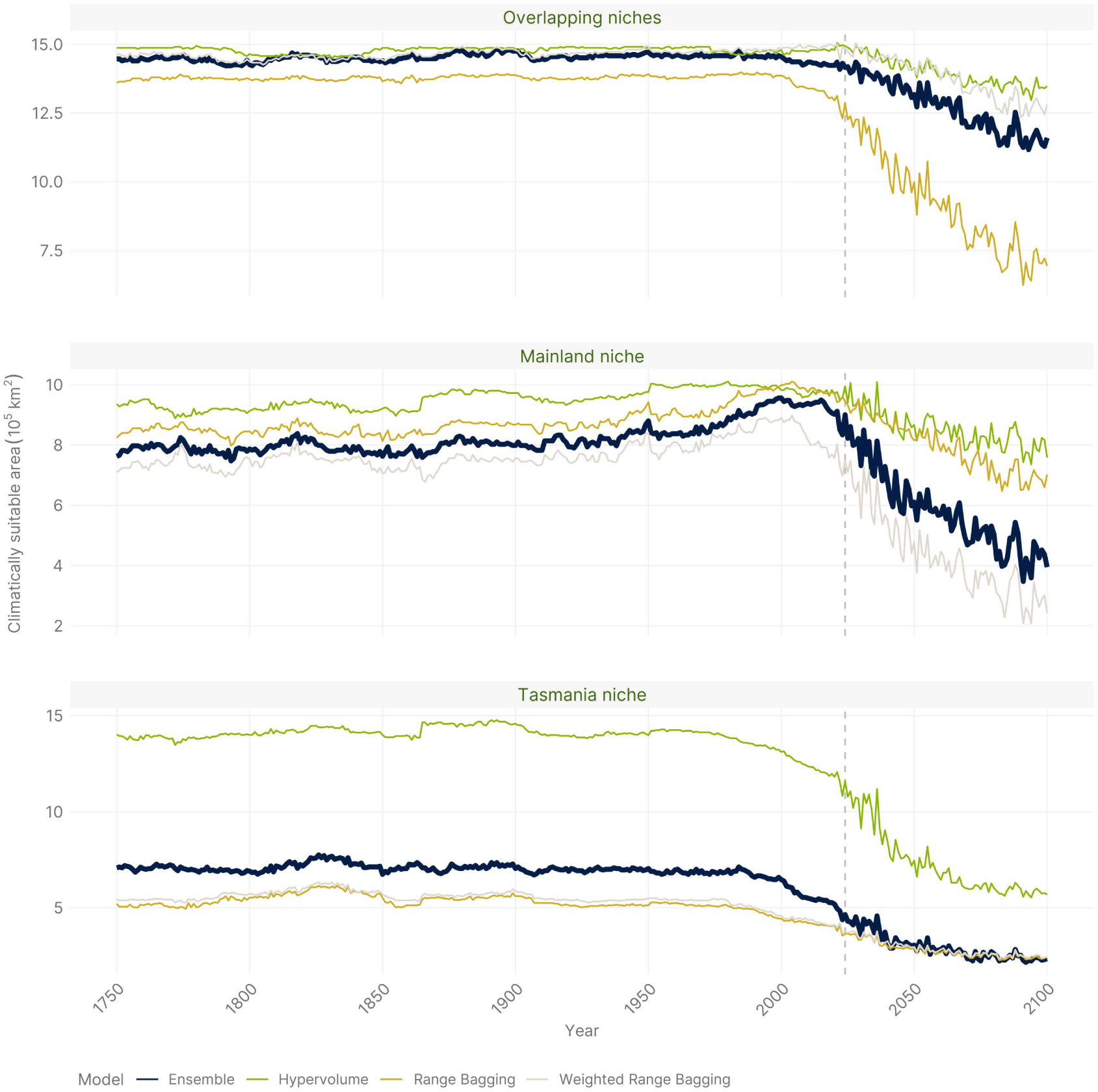
Sum of suitable climate area through time, defined as all cells with climate suitability above the threshold value for *Perameles gunnii* across all scenarios and climate suitability models. The vertical dashed lines represent the year 2024. Note the y axes have different scales in each panel.

#### Parameter estimates of population dynamics leading to extirpation from the mainland

Our SEPM framework managed to recreate the extirpation pattern of both *D. viverrinus* and *P. gunnii* from mainland Australia, with populations crashing rapidly in the second half of the 19^th^ century following the spread of invasive foxes and increasing harvest rates (Fig. S12).

**Figure S12.**
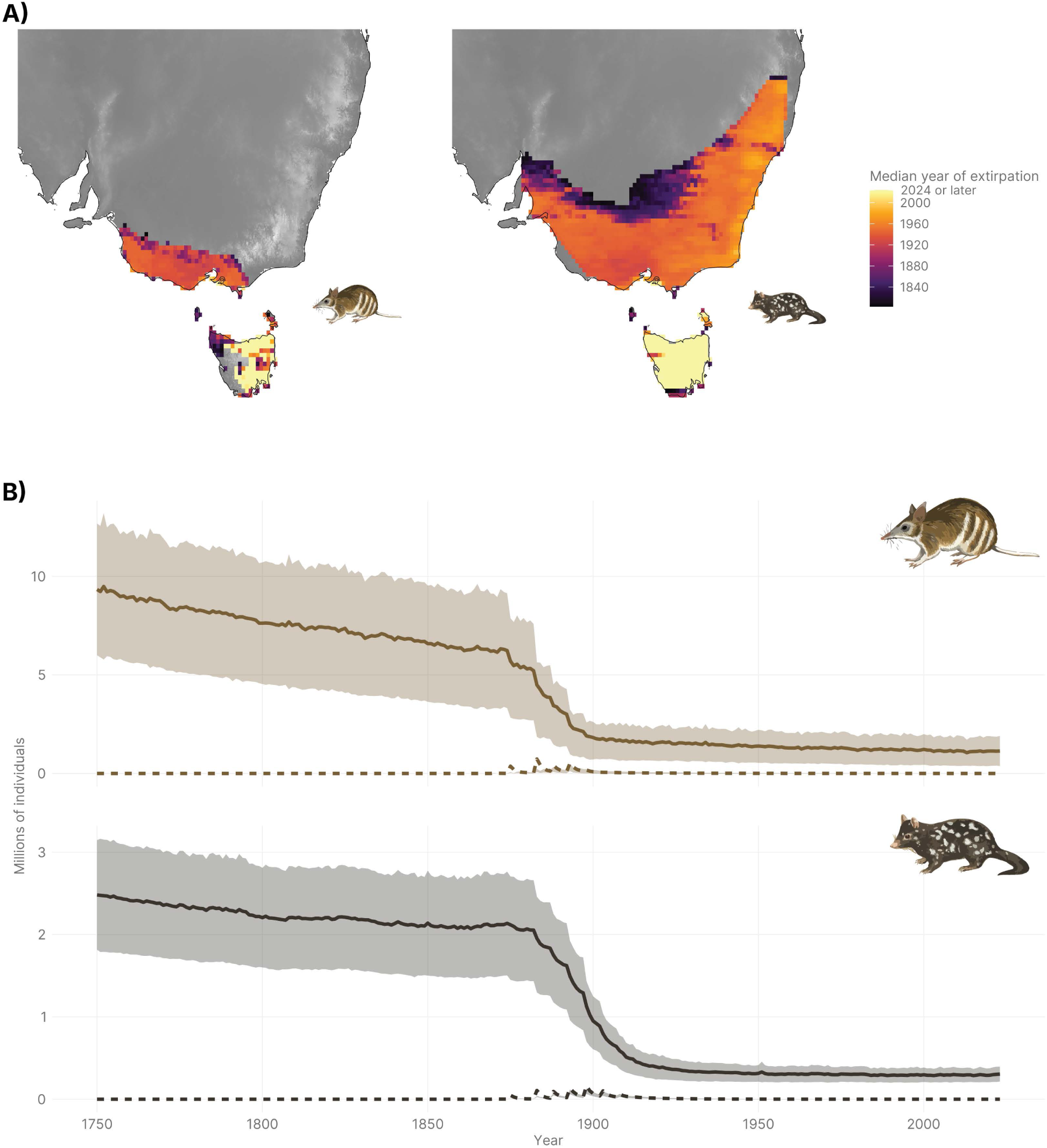
A) Maps showing median year of extirpation for *Perameles gunnii* (left) and *Dasyurus viverrinus* (right); B) Predicted population size (continuous line) and predation by foxes (dashed line) through time (1750–2024) across the study area for *Perameles gunnii* (top) and *Dasyurus viverrinus* (bottom). Both measures are calculated as the sum across the entire study area of the mean abundance and harvest across the 0.025% simulations closest to the validation targets, weighted by the inverse of the Euclidean distance of each simulation to the targets. Median year of extirpation is calculated for each cell across the same subset of simulations and using the same weighting scheme. Shaded areas represent 95% confidence intervals.

However, a decline in populations was evident even before the introduction of foxes, likely due to a decline in climatic suitability particularly in the lowland areas of Victoria and southern South Australia (Fig. 1 in main text).

The posterior parameter estimates suggest that both species’ populations are highly fluctuating (modal *standard_deviation* = 0.49 and 0.64 for *D. viverrinus* and *P. gunnii*, respectively), but *P. gunnii* achieves a faster growth rate compared to *D. viverrinus* (modal *growth_rate_max* = 0.15 and 0.48 for *D. viverrinus* and *P. gunnii*, respectively), which is expected given adults of the latter are larger and in mammals population growth scales negatively with body size (Thompson 1987). Both species have short gestation periods and large litters (Dufty 1991, 1994; Backhouse *et al*. 1994; Jones & Rose 2001; Winnard & Coulson 2008), but *P. gunnii* females are able to produce multiple litters per year, whereas *D. viverrinus* females typically breed only once per year, in winter, unless conception fails or the litter is lost early in the season (Fletcher 1985). Seasonal population fluctuations due to juvenile mortality (Godsell 1983), coupled with a short life span, leads to high population turnover in *D. viverrinus* (Jones & Rose 2001). Fluctuating population growth rates coupled with a relatively high threshold for local population extirpation, at least for *D. viverrinus* (modal *abundance_threshold* = 0.007 individuals / ha, roughly 7.2% of the maximum population density under ideal conditions), could make species quite vulnerable to extinction, which may explain why our reconstructions showed mild but consistent population declines even before the introduction of foxes (Fig. S12). This trend suggests that mainland populations would have diminished, and some would have perhaps experienced local extirpation, even without pressure from introduced predators and due to changing climates alone. However, our models suggest that introduced foxes provided the death knell for Australian mainland populations due to extremely high predation rates (modal *harvest_max* = 0.784 and 0.715 for *D. viverrinus* and *P. gunnii*, respectively), coupled with low values of the *z* parameter (modal *harvest_z* = 1.12 and 1.23 for *D. viverrinus* and *P. gunnii*, respectively).

Predation is modelled here using an equilibrium predation pressure functional response (Alroy 2001; Brook & Johnson 2006; Tomlinson *et al*. 2023). When *z* is equal to 1, the function is monotonic (type II interaction), meaning predation is dependent only on prey density and predator satiation (Alroy 2001), a scenario likely to lead to extirpation due to high predation pressure at low prey abundances. At values of *z* above 1.5, predation becomes increasingly density-dependent (type III interaction), whereby predation rate is lower at lower prey density—*i.e.*, predators are less successful at capturing prey when prey is less abundant. Our estimated values of *z* suggest that for *D. viverrinus* and *P. gunnii* this is not the case, and predation rate more closely follows a type II interaction, implying prey naivety (Sinclair *et al*. 1998). As a direct outcome of this and the high estimated maximum harvest rates, the population-level harvest rate (*i.e.*, the proportion of the population that is killed due to predation) is at its highest—and much greater than over 50%—when population density is low (Fig. S13), either due to a lower carrying capacity or because predation or stochastic events lead to declines. This could explain why populations on the mainland deteriorated so quickly after fox introduction—a decline which is also supported by genetic evidence from the last remaining wild mainland population of *P. gunnii* in Hamiton, Victoria in 1992 (Weeks *et al*. 2013).

**Figure S13.**
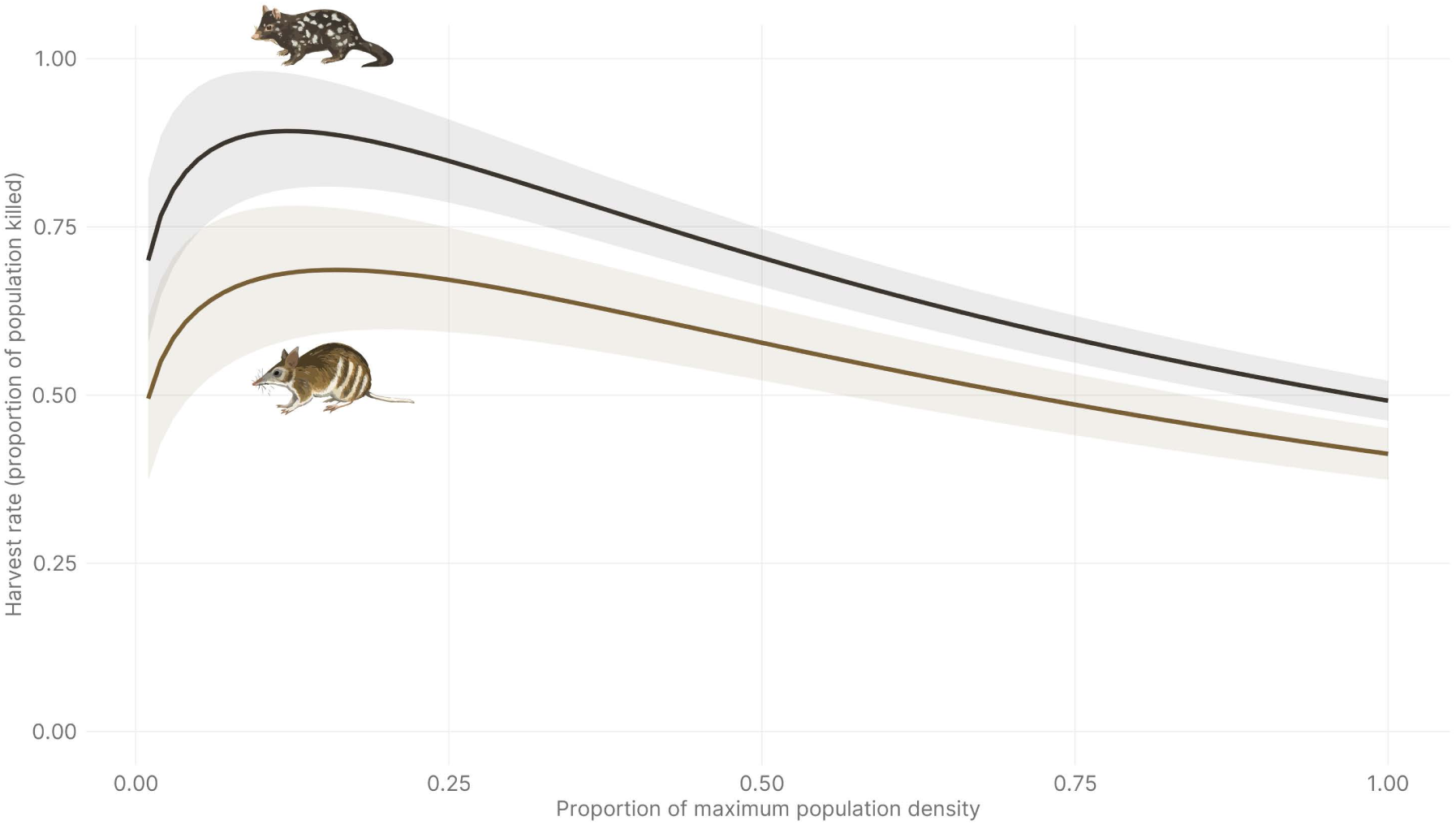
Population level harvest rates as a function of relative population density (out of the maximum population density) for *Perameles gunnii* and *Dasyurus viverrinus*, calculated from the posterior estimates for maximal predation rate and the z parameter.

It is important to note that our models may have underestimated population growth dynamics, and/or overestimated predation rates and abundance thresholds (Tomlinson *et al*. 2023). The reasons for this are tow-fold: first, while we modelled predation by foxes alone, which are considered to be the main reason for our target species’ declines (Jones & Rose 2001; Kinnear *et al*. 2002; DSE Victoria 2009), predation by introduced cats (*Felis catus*) could also have contributed to the decline of these two species (Brown 1989; Seebeck *et al*. 1990; Kinnear *et al*. 2002; Radford *et al*. 2018). Cats and foxes could also interact with each other, and both be suppressed by the presence of dingoes (*Canis familiaris dingo*), an apex predator in many Australian terrestrial habitats (Johnson & VanDerWal 2009; Brook *et al*. 2012; Fancourt *et al*. 2019; Stobo-Wilson *et al*. 2022). Second, the decline of *P. gunnii* from Victoria’s south-west is also closely tied to the widespread conversion of native grasslands and grassy woodlands in the volcanic plains to agricultural landscapes (Brown 1989; Seebeck *et al*. 1990). Optimal habitat for *P. gunnii* is defined by high structural complexity and habitat heterogeneity (Dufty 1994), meaning that homogenous agricultural fields with low complexity can be unsuitable even if climatically the area is within the species’ niche. A modelling framework that incorporates all these interacting processes will be able to give much more accurate estimates of population dynamics and allow for a careful exploration of the leading causes of *D. viverrinus* and *P. gunnii* decline.

#### Implications of niche divergence between mainland and Tasmanian populations

We ran all simulations under two scenarios—‘Overlapping niches’ and ‘Divergent niches’. We found that under the ‘Divergent niches’ scenario, climatic suitability trends and population trajectories vary widely between mainland and Tasmanian populations (Figs. S7–S10) due to low overlap particularly in thermal maxima between the two (Fig. S11). This could have important implications for reintroduction efforts—for example, for both species current sanctuary populations are projected to collapse rapidly if they are comprised of animals with a climatically divergent Tasmanian niche (Figs. S14–S15).

**Figure S11.**
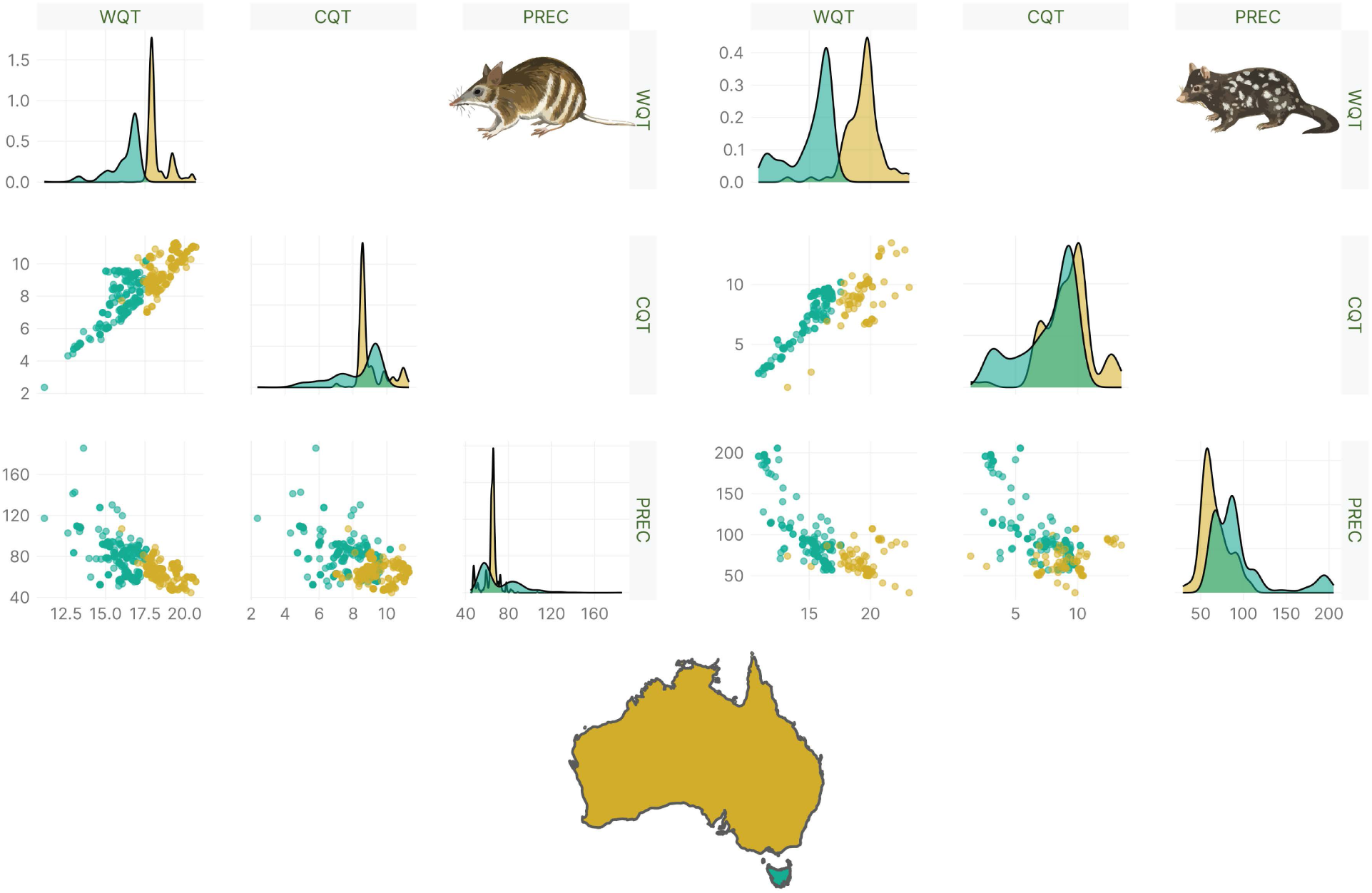
Distributions of WQT (average monthly temperatures for the warmest quarter), CQT (average monthly temperatures for the coldest quarter) and PREC (total annual rainfall) for occurrence records of *Perameles gunnii* (left) and *Dasyurus viverrinus* (right) from mainland Australia (orange) and Tasmania (light blue). The bottom triangle shows scatterplots with the variable in the corresponding column on the x axis and the variable in the corresponding row on the y axis. The diagonal shows univariate density plots.

The mainland and Tasmanian populations of *P. gunnii* are strongly genetically differentiated (Robinson *et al*. 1993; Robinson 1995; Weeks *et al*. 2013), to the point where they are considered by some to be distinct subspecies. Sanctuary populations of *P. gunnii* on the mainland are derived from a captive-bred colony originally sourced from the last remnant mainland population in Hamilton, Victoria (Weeks *et al*. 2013). In part due to the small founder populations (Myorniuk 1993), these reintroduced sanctuary populations have experienced rapid declines in genetic diversity over the last decades (Weeks *et al*. 2013). Loss of genetic diversity is likely to increase the risk of extinction and reduce adaptive potential (Willi *et al*. 2006; Markert *et al*. 2010; Weeks *et al*. 2011), and small captive-bred populations could be vulnerable to the accumulation of deleterious alleles (Agrawal & Whitlock 2012). To combat this, ‘genetic rescue’ (Hedrick 1995; Hedrick & Fredrickson 2010; Weeks *et al*. 2011) has been suggested as a management solution by introducing individuals from Tasmania to the sanctuary populations in order to bolster genetic diversity. However, a possible risk in a ‘genetic rescue’ approach is if the source and recipient populations are locally adapted to different conditions (Weeks *et al*. 2011). Our simulations suggest that, if the fundamental niches of Tasmanian and mainland populations are fully diverged, then Tasmanian animals are already maladapted to persist in mainland sites. Therefore, a careful assessment of niche divergence between Tasmanian and mainland *P. gunnii* should be undertaken before ‘genetic rescue’ is implemented.

Conversely, the original mainland populations of *D. viverrinus* are completely extinct—they disappeared much earlier than *P. gunnii* (1960s vs 1990s) and reintroduced mainland populations are derived solely from translocated Tasmanian animals (Woinarski *et al*. 2014; Wilson *et al*. 2020, 2023). Little is known about the genetics of reintroduced *D. viverrinus* populations in sanctuaries. However, there is significant population structure among Tasmanian populations of the species (Firestone *et al*. 2000; Cardoso *et al*. 2014), with reduced genetic diversity in coastal, peripheral populations (Cardoso *et al*. 2014). These are also the regions where our models predict declines in climatic suitability (Fig. 1A in main text). For *D. viverrinus*, since all current mainland populations are derived from translocated Tasmanian individuals, if niche divergence has occurred then maintaining evolutionary resilience to climate change will be extremely challenging. However, current population densities of *D. viverrinus* at Mulligan’s Flat (∼0.1 individuals/ha; Wilson *et al*., 2023) are more consistent with our predictions under an ‘Overlapping niches’ scenario (95% CI 0–0.08 individuals/ha; Fig. S14) rather than the much lower predictions under the ‘Tasmania niche’ scenario (95% CI 0– 0.05 individuals/ha), suggesting niches have not diverged substantially in this species.

**Figure S14.**
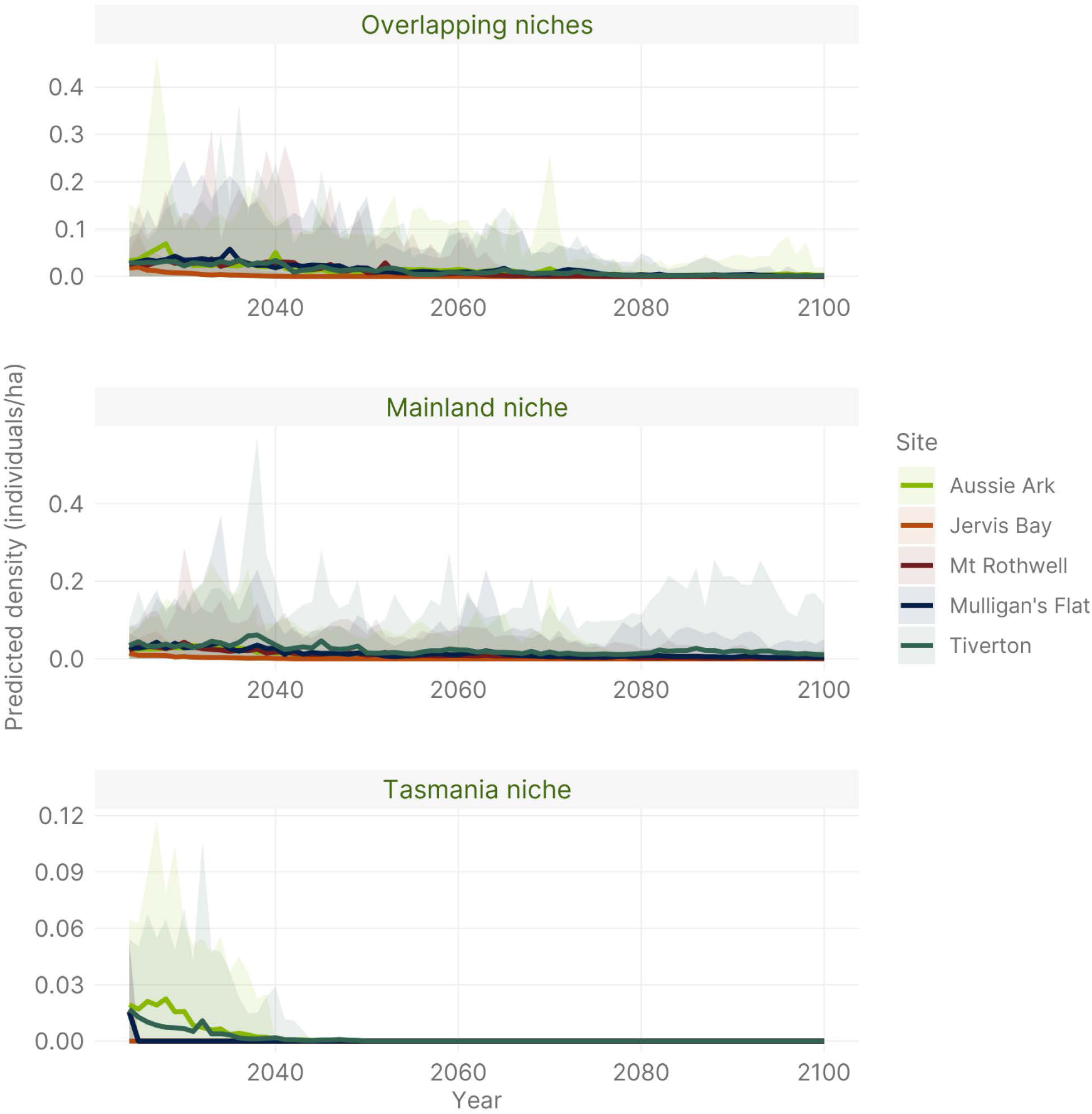
Predicted population trajectories of reintroduced mainland populations of *Dasyurus viverrinus* in predator-free exclosures under different scenarios of modelled habitat suitability. The lines show the mean population densities from all 50 simulations, and the shaded areas show the 95% CI around the mean.

**Figure S15.**
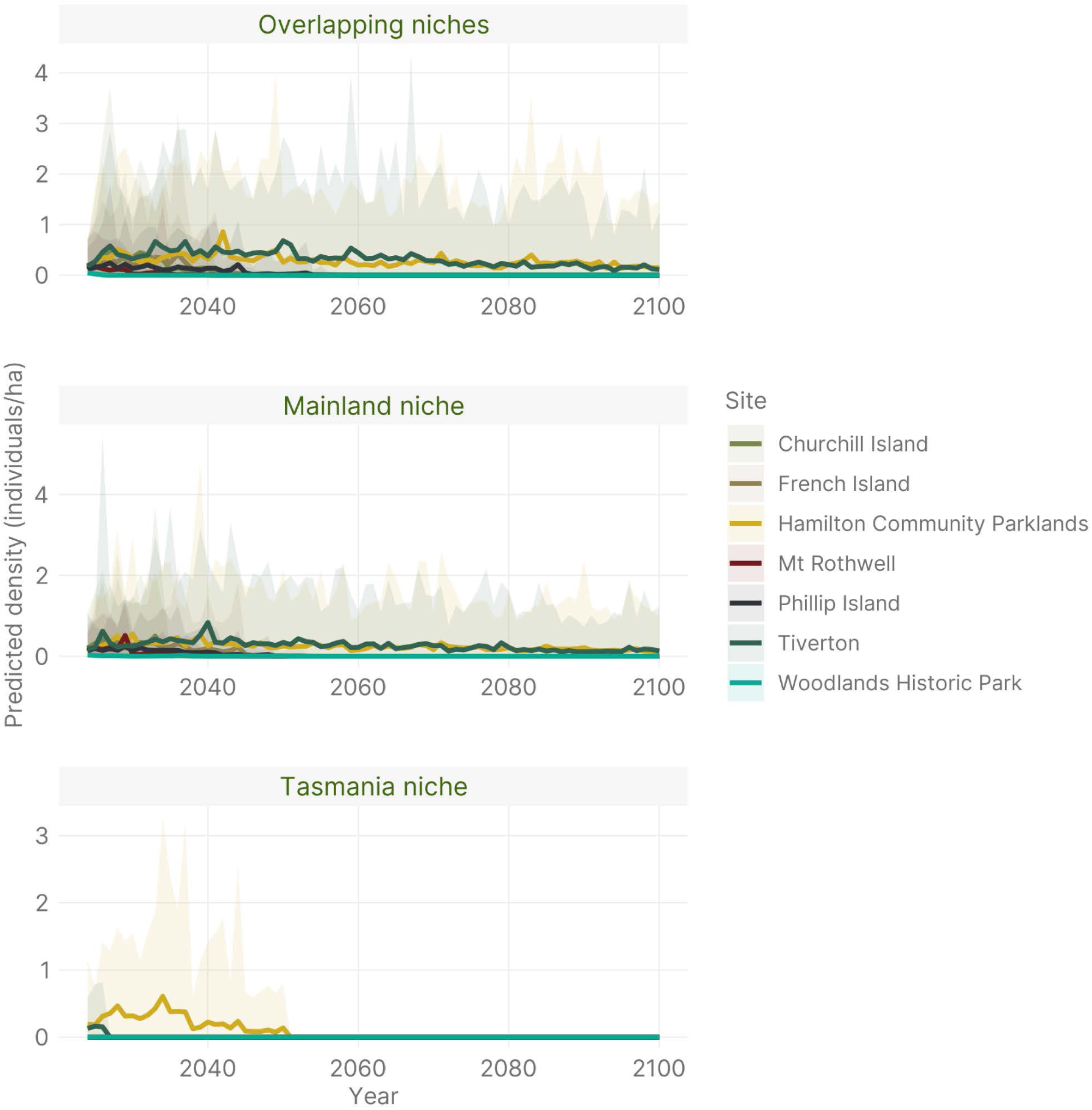
Predicted population trajectories for reintroduced mainland populations of *Perameles gunnii* in predator-free exclosures under different scenarios of modelled habitat suitability. The lines show the mean population densities from all 50 simulations, and the shaded areas show the 95% CI around the mean.

## References

Agrawal, A.F. & Whitlock, M.C. (2012). Mutation load: the fitness of individuals in populations where deleterious alleles are abundant. Annu Rev Ecol Evol Syst, 43, 115–135.

Alroy, J. (2001). A multispecies overkill simulation of the end-Pleistocene megafaunal mass extinction. Science (1979), 292, 1893–1896.

Araújo, M.B., Pearson, R.G., Thuiller, W. & Erhard, M. (2005). Validation of species-climate impact models under climate change. Glob Chang Biol, 11, 1504–1513.

Backhouse, G.N., Clark, T.W. & Reading, R.P. (1994). Reintroductions for recovery of the eastern barred bandicoot Perameles gunnii in Victoria, Australia. In: *Reintroduction Biology of Australian and New Zealand Fauna* (ed. Serena, M.). Surrey Beatty & Sons, Cipping Norton, NSW, pp. 209–218.

Barghi, N., Hermisson, J. & Schlötterer, C. (2020). Polygenic adaptation: a unifying framework to understand positive selection. Nat Rev Genet, 21, 769–781.

Barrett, R.D. & Schluter, D. (2008). Adaptation from standing genetic variation. Trends Ecol Evol, 23, 38–44.

Belbin, L., Wallis, E., Hobern, D. & Zerger, A. (2021). The Atlas of Living Australia: History, current state and future directions. Biodivers Data J, 9, e65023.

Beyer, R., Krapp, M. & Manica, A. (2020). An empirical evaluation of bias correction methods for palaeoclimate simulations. Climate of the Past, 16, 1493–1508.

Blonder, B., Lamanna, C., Violle, C. & Enquist, B.J. (2014). The n-dimensional hypervolume. Global Ecology and Biogeography, 23, 595–609.

Blonder, B., Morrow, C.B., Harris, D.J., Brown, S., Butruille, G., Laini, A. & Chen, D. (2023). hypervolume: High Dimensional Geometry, Set Operations, Projection, and Inference Using Kernel Density Estimation, Support Vector Machines, and Convex Hulls.

Bozinovic, F. & Pörtner, H.-O. (2015). Physiological ecology meets climate change. Ecol Evol, 5, 1025–1030.

Brook, B.W. & Johnson, C.N. (2006). Selective hunting of juveniles as a cause of the imperceptible overkill of the Australian Pleistocene megafauna. Alcheringa: an Australasian Journal of Palaeontology, 30, 39–48.

Brook, L.A., Johnson, C.N. & Ritchie, E.G. (2012). Effects of predator control on behaviour of an apex predator and indirect consequences for mesopredator suppression. Journal of Applied Ecology, 49, 1278–1286.

Brown, P.R. (1989). *Management plan for the conservation of the eastern barred bandicoot*, Perameles gunnii, *in Victoria: A report to the Department of Conservation, Forests, and Lands*, Victoria and the World Wildlife Fund Australia.

Brown, S.C., Wigley, T.M.L., Otto-Bliesner, B.L. & Fordham, D.A. (2020). *StableClim*, continuous projections of climate stability from 21000 BP to 2100 CE at multiple spatial scales. Sci Data, 7, 335.

Cardoso, M.J., Mooney, N., Eldridge, M.D.B., Firestone, K.B. & Sherwin, W.B. (2014). Genetic monitoring reveals significant population structure in eastern quolls: implications for the conservation of a threatened carnivorous marsupial. Aust Mammal, 36, 169–177.

Csilléry, K., François, O. & Blum, M.G.B. (2012). abc: an R package for approximate Bayesian computation (ABC). Methods Ecol Evol, 3, 475–479.

Drake, J.M. (2015). Range bagging: a new method for ecological niche modelling from presence-only data. J R Soc Interface, 12, 20150086.

DSE Victoria. (2009). *Eastern barred bandicoot (mainland)* Perameles gunni *(unnamed subspecies)*. Action statement no. 4 (revised 2009).

Dufty, A.C. (1991). Some population characteristics of *Perameles gunnii* in Victoria. Wildlife Research, 18, 355–365.

Dufty, A.C. (1994). Habitat and spatial requirements of the eastern barred bandicoot (*Perameles gunnii*) at Hamilton, Victoria. Wildlife Research, 21, 459–471.

Evans, M.J., Batson, W.G., Gordon, I.J., Belton, E., Chaseling, T., Fletcher, D., Harrison, M., McElroy, T., Mungoven, A., Newport, J., Pierson, J., Portas, T., Swain, S., Wimpenny, C. & Manning, A.D. (2021). The ‘Goldilocks Zone’ of predation: the level of fox control needed to select predator resistance in a reintroduced mammal in Australia. Biodivers Conserv, 30, 1731–1752.

Evans, M.J., Weeks, A.R., Scheele, B.C., Gordon, I.J., Neaves, L.E., Andrewartha, T.A., Brockett, B., Rapley, S., Smith, K.J., Wilson, B.A. & Manning, A.D. (2022). Coexistence conservation: Reconciling threatened species and invasive predators through adaptive ecological and evolutionary approaches. Conserv Sci Pract, 4.

Fairfax, R.J. (2019). Dispersal of the introduced red fox (*Vulpes vulpes*) across Australia. Biol Invasions, 21, 1259–1268.

Fancourt, B.A., Cremasco, P., Wilson, C. & Gentle, M.N. (2019). Do introduced apex predators suppress introduced mesopredators? A multiscale spatiotemporal study of dingoes and feral cats in Australia suggests not. Journal of Applied Ecology, 56, 2584– 2595.

Firestone, K.B., Houlden, B.A., Sherwin, W.B. & Geffen, E. (2000). Variability and differentiation of microsatellites in the genus *Dasyurus* and conservation implications for the large Australian carnivorous marsupials. Conservation Genetics, 1, 115–133.

Fletcher, T.P. (1985). Aspects of reproduction in the male eastern quoll, *Dasyurus viverrinus* (Shaw) (Marsupialia: Dasyuridae), with notes on polyestry in the female. Aust J Zool, 33, 101–110.

Fordham, D.A., Akçakaya, H.R., Araújo, M.B., Keith, D.A. & Brook, B.W. (2013a). Tools for integrating range change, extinction risk and climate change information into conservation management. Ecography, 36, 956–964.

Fordham, D.A., Akçakaya, H.R., Brook, B.W., Rodríguez, A., Alves, P.C., Civantos, E., Triviño, M., Watts, M.J. & Araújo, M.B. (2013b). Adapted conservation measures are required to save the Iberian lynx in a changing climate. Nat Clim Chang, 3, 899–903.

Fordham, D.A., Haythorne, S., Brown, S.C., Buettel, J.C. & Brook, B.W. (2021). poems: R package for simulating species’ range dynamics using pattern-oriented validation. Methods Ecol Evol, 12, 2364–2371.

Fordham, D.A., Saltré, F., Haythorne, S., Wigley, T.M.L., Otto-Bliesner, B.L., Chan, K.C. & Brook, B.W. (2017). PaleoView: a tool for generating continuous climate projections spanning the last 21000 years at regional and global scales. Ecography, 40, 1348–1358.

Frankham, R., Bradshaw, C.J.A. & Brook, B.W. (2014). Genetics in conservation management: Revised recommendations for the 50/500 rules, Red List criteria and population viability analyses. Biol Conserv, 170, 56–63.

Franklin, I.R. (1980). Evolutionary change in small populations. In: Conservation biology: an evolutionary-ecological perspective (eds. Soulé, M.E. & Wilcox, B.A.). Sinauer Associates, Inc., Sunderland, Massachusetts, pp. 135–149.

Franklin, I.R. & Frankham, R. (1998). How large must populations be to retain evolutionary potential? Anim Conserv, 1, 69–70.

Godsell, J. (1983). *Ecology of the eastern quoll* Dasyurus viverrinus, (Dasyuridae: Marsuipalia) (PhD).

Grose, M., Abbs, D., Bhend, J., Chiew, F., Church, J., Ekström, M., Kirono, D., Lenton, A., Lucas, C., McInnes, K., Moise, A., Monselesan, D., Mpelasoka, F., Webb, L. & Whetton, P. (2015). Southern Slopes Cluster Report, Climate Change in Australia Projections for Australia’s Natural Resource Management Regions: Cluster Reports. In: (eds. Ekström, M., Whetton, P., Gerbing, C., Grose, M., Webb, L. & Risbey, J.). CSIRO and Bureau of Meteorology, Australia.

Guzella, T.S., Dey, S., Chelo, I.M., Pino-Querido, A., Pereira, V.F., Proulx, S.R. & Teotónio, H. (2018). Slower environmental change hinders adaptation from standing genetic variation. PLoS Genet, 14, e1007731.

Habel, K., Grasman, R., Gramacy, R., Mozharovskyi, P. & Sterratt, D. (2022). geometry: Mesh Generation and Surface Tessellation.

Haller, B.C. & Messer, P.W. (2023). SLiM 4: Multispecies eco-evolutionary modeling. Am Nat, 201, E127–E139.

Haythorne, S., Pilowsky, J., Brown, S. & Fordham, D. (2024). paleopop: Pattern-Oriented Modeling Framework for Coupled Niche-Population Paleo-Climatic Models.

Hayward, L.K. & Sella, G. (2022). Polygenic adaptation after a sudden change in environment. Elife, 11.

Hayward, M.W. & Kerley, G.I.H. (2009). Fencing for conservation: Restriction of evolutionary potential or a riposte to threatening processes? Biol Conserv, 142, 1–13.

Hayward, M.W., Moseby, K. & Read, J.L. (2014). The role of predator exclosures in the conservation of Australian fauna. In: Carnivores of Australia: Past, Present and Future (eds. Glen, A. & Dickman, C.). CSIRO Publishing, Collingwood, Australia, pp. 353–371.

Hedrick, P.W. (1995). Gene flow and genetic restoration: the Florida panther as a case study. Conservation Biology, 9, 996–1007.

Hedrick, P.W. & Fredrickson, R. (2010). Genetic rescue guidelines with examples from Mexican wolves and Florida panthers. Conservation Genetics, 11, 615–626.

Hermisson, J. & Pennings, P.S. (2005). Soft sweeps: molecular population genetics of adaptation from standing genetic variation. Genetics, 169, 2335–2352.

Hirzel, A.H., Le Lay, G., Helfer, V., Randin, C. & Guisan, A. (2006). Evaluating the ability of habitat suitability models to predict species presences. Ecol Modell, 199, 142–152.

Hoffmann, A.A. & Sgrò, C.M. (2011). Climate change and evolutionary adaptation. Nature, 470, 479–485.

Hogg, C.J., Edwards, R.J., Farquharson, K.A., Silver, L.W., Brandies, P., Peel, E., Escalona, M., Jaya, F.R., Thavornkanlapachai, R., Batley, K., Bradford, T.M., Chang, J.K., Chen, Z., Deshpande, N., Dziminski, M., Ewart, K.M., Griffith, O.W., Marin Gual, L., Moon, K.L., Travouillon, K.J., Waters, P., Whittington, C.M., Wilkins, M.R., Helgen, K.M., Lo, N., Ho, S.Y.W., Ruiz Herrera, A., Paltridge, R., Marshall Graves, J.A., Renfree, M., Shapiro, B., Ottewell, K., Gibson, C., Maxwell, R., Spencer, Z., Napangati, Y., Butler, M., West, J., West, J., James, M., Napangati, N., Gibson, L., West, P., Gibson, A., West, S., West, K., Japaltjari, W., Blackwood, E., Rachel Paltridge & Belov, K. (2024). Extant and extinct bilby genomes combined with Indigenous knowledge improve conservation of a unique Australian marsupial. Nat Ecol Evol, 8, 1311–1326.

Innes, J., Burns, B., Sanders, A. & Hayward, M.W. (2015). The impact of private sanctuary networks on reintroduction programs. In: Advances in Reintroduction Biology of Australian and New Zealand Fauna (eds. Armstrong, D., Hayward, M., Moro, D. & Seddon, P.). CSIRO Publishing, Victoria, Australia, pp. 185–199.

Johnson, C.N. (2006). Australia’s mammal extinctions: a 50,000-year history. Cambridge University Press, Cambridge, UK.

Johnson, C.N. & VanDerWal, J. (2009). Evidence that dingoes limit abundance of a mesopredator in eastern Australian forests. Journal of Applied Ecology, 46, 641–646.

Jones, M.E. & Rose, R.K. (2001). Dasyurus viverrinus. Mammalian Species, 677, 1–9.

Keith, D.A., Akçakaya, H.R., Thuiller, W., Midgley, G.F., Pearson, R.G., Phillips, S.J., Regan, H.M., Araújo, M.B. & Rebelo, T.G. (2008). Predicting extinction risks under climate change: coupling stochastic population models with dynamic bioclimatic habitat models. Biol Lett, 4, 560–563.

Kinnear, J.E., Sumner, N.R. & Onus, M.L. (2002). The red fox in Australia—an exotic predator turned biocontrol agent. Biol Conserv, 108, 335–359.

Lacy, R.C. (1987). Loss of genetic diversity from managed populations: interacting effects of drift, mutation, immigration, selection, and population subdivision. Conservation Biology, 1, 143–158.

Lai, Y.-T., Yeung, C.K.L., Omland, K.E., Pang, E.-L., Hao, Y., Liao, B.-Y., Cao, H.-F., Zhang, B.-W., Yeh, C.-F., Hung, C.-M., Hung, H.-Y., Yang, M.-Y., Liang, W., Hsu, Y.-C., Yao, C.-T., Dong, L., Lin, K. & Li, S.-H. (2019). Standing genetic variation as the predominant source for adaptation of a songbird. Proceedings of the National Academy of Sciences, 116, 2152–2157.

Lande, R. (1995). Mutation and Conservation. Conservation Biology, 9, 782–791.

Legge, S., Woinarski, J.C.Z., Burbidge, A.A., Palmer, R., Ringma, J., Radford, J.Q., Mitchell, N., Bode, M., Wintle, B., Baseler, M., Bentley, J., Copley, P., Dexter, N., Dickman, C.R., Gillespie, G.R., Hill, B., Johnson, C.N., Latch, P., Letnic, M., Manning, A., McCreless, E.E., Menkhorst, P., Morris, K., Moseby, K., Page, M., Pannell, D. & Tuft, K. (2018). Havens for threatened Australian mammals: the contributions of fenced areas and offshore islands to the protection of mammal species susceptible to introduced predators. Wildlife Research, 45, 627.

Mace, G.M. & Purvis, A. (2008). Evolutionary biology and practical conservation: bridging a widening gap. Mol Ecol, 17, 9–19.

Markert, J.A., Champlin, D.M., Gutjahr-Gobell, R., Grear, J.S., Kuhn, A., McGreevy Jr, T.J., Roth, A., Bagley, M.J. & Nacci, D.E. (2010). Population genetic diversity and fitness in multiple environments. BMC Evol Biol, 10, 1–13.

Mee, J.A. & Yeaman, S. (2019). Unpacking conditional neutrality: genomic signatures of selection on conditionally beneficial and conditionally deleterious mutations. Am Nat, 194, 529–540.

Murali, G., Iwamura, T., Meiri, S. & Roll, U. (2023). Future temperature extremes threaten land vertebrates. Nature, 615, 461–467.

Myorniuk, P. (1993). Eastern barred bandicoot recovery. Newsletter no. 3.

Nogués-Bravo, D. (2009). Predicting the past distribution of species climatic niches. Global Ecology and Biogeography, 18, 521–531.

Orr, H.A. (2010). The population genetics of beneficial mutations. Philosophical Transactions of the Royal Society B: Biological Sciences, 365, 1195–1201.

Pacifici, M., Foden, W.B., Visconti, P., Watson, J.E.M., Butchart, S.H.M., Kovacs, K.M., Scheffers, B.R., Hole, D.G., Martin, T.G., Akçakaya, H.R., Corlett, R.T., Huntley, B., Bickford, D., Carr, J.A., Hoffmann, A.A., Midgley, G.F., Pearce-Kelly, P., Pearson, R.G., Williams, S.E., Willis, S.G., Young, B. & Rondinini, C. (2015). Assessing species vulnerability to climate change. Nat Clim Chang, 5, 215–224.

Pateiro-Lopez, B. & Rodriguez-Casal, A. (2022). alphahull: Generalization of the Convex Hull of a Sample of Points in the Plane.

Pearson, R.G. & Dawson, T.P. (2003). Predicting the impacts of climate change on the distribution of species: are bioclimate envelope models useful? Global Ecology and Biogeography, 12, 361–371.

Pritchard, J.K. & Di Rienzo, A. (2010). Adaptation – not by sweeps alone. Nat Rev Genet, 11, 665–667.

Prowse, T.A.A., Bradshaw, C.J.A., Delean, S., Cassey, P., Lacy, R.C., Wells, K., Aiello-Lammens, M.E., Akçakaya, H.R. & Brook, B.W. (2016). An efficient protocol for the global sensitivity analysis of stochastic ecological models. Ecosphere, 7, e01238.

R Core Team. (2022). R: a language and environment for statistical computing.

R Core Team. (2025). R: a language and environment for statistical computing.

Radford, J.Q., Woinarski, J.C.Z., Legge, S., Baseler, M., Bentley, J., Burbidge, A.A., Bode, M., Copley, P., Dexter, N., Dickman, C.R., Gillespie, G., Hill, B., Johnson, C.N., Kanowski, J., Latch, P., Letnic, M., Manning, A., Menkhorst, P., Mitchell, N., Morris, K., Moseby, K., Page, M. & Ringma, J. (2018). Degrees of population-level susceptibility of Australian terrestrial non-volant mammal species to predation by the introduced red fox (*Vulpes vulpes*) and feral cat (*Felis catus*). Wildlife Research, 45, 645–657.

Ricker, W.E. (1954). Stock and recruitment. Journal of the Fisheries Board of Canada, 11, 559– 623.

Ringma, J., Legge, S., Woinarski, J., Radford, J., Wintle, B. & Bode, M. (2018). Australia’s mammal fauna requires a strategic and enhanced network of predator-free havens. Nat Ecol Evol, 2, 410–411.

Robinson, N.A. (1995). Implications from mitochondrial DNA for management to conserve the eastern barred bandicoot (*Perameles gunnii*). Conservation Biology, 9, 114–125.

Robinson, N.A., Murray, N.D. & Sherwin, W.B. (1993). VNTR loci reveal differentiation between and structure within populations of the eastern barred bandicoot *Perameles gunnii*. Mol Ecol, 2, 195–207.

Seebeck, J.H., Bennett, A.F. & Dufty, A.C. (1990). Status, distribution, and biogeography of the eastern barred bandicoot, Perameles gunnii in Victoria. In: Management and Conservation of Small Populations (eds. Clark, T.W. & Seebeck, J.H.). Chicago Zoological Society, Brookfield, Illinois, pp. 21–32.

Sgrò, C.M., Lowe, A.J. & Hoffmann, A.A. (2011). Building evolutionary resilience for conserving biodiversity under climate change. Evol Appl, 4, 326–337.

Shaw, R.E., Farquharson, K.A., Bruford, M.W., Coates, D.J., Elliott, C.P., Mergeay, J., Ottewell, K.M., Segelbacher, G., Hoban, S., Hvilsom, C., Pérez-Espona, S., Ruņģis, D., Aravanopoulos, F., Bertola, L.D., Cotrim, H., Cox, K., Cubric-Curik, V., Ekblom, R., Godoy, J.A., Konopiński, M.K., Laikre, L., Russo, I.-R.M., Veličković, N., Vergeer, P., Vilà, C., Brajkovic, V., Field, D.L., Goodall-Copestake, W.P., Hailer, F., Hopley, T., Zachos, F.E., Alves, P.C., Biedrzycka, A., Binks, R.M., Buiteveld, J., Buzan, E., Byrne, M., Huntley, B., Iacolina, L., Keehnen, N.L.P., Klinga, P., Kopatz, A., Kurland, S., Leonard, J.A., Manfrin, C., Marchesini, A., Millar, M.A., Orozco-terWengel, P., Ottenburghs, J., Posledovich, D., Spencer, P.B., Tourvas, N., Unuk Nahberger, T., van Hooft, P., Verbylaite, R., Vernesi, C. & Grueber, C.E. (2025). Global meta-analysis shows action is needed to halt genetic diversity loss. Nature, 638, 704–710.

Silverman, B.W. (1986). Density estimation for statistics and data analysis. Chapman & Hall, London, UK.

Sinclair, A.R.E., Pech, R.P., Dickman, C.R., Hik, D., Mahon, P. & E, N. (1998). Predicting effects of predation on conservation of endangered prey. Conservation Biology, 12, 564– 575.

Stephens, P.A. & Sutherland, W.J. (1999). Consequences of the Allee effect for behaviour, ecology and conservation. Trends Ecol Evol, 14, 401–405.

Stephens, P.A., Sutherland, W.J. & Freckleton, R.P. (1999). What is the Allee effect? Oikos, 87, 185–190.

Stobo-Wilson, A., Murphy, B.P., Legge, S.M., Caceres-Escobar, H., Chapple, D.G., Crawford, H.M., Dawson, S.J., Dickman, C.R., Doherty, T.S., Fleming, P.A., Garnett, S.T., Gentle, M., Newsome, T.M., Palmer, R., Rees, M.W., Ritchie, E.G., Speed, J., Stuart, J.-M., Suarez-Castro, A.F., Thompson, E., Tulloch, A., Turpin, J.M. & Woinarski, J.C.Z. (2022). Counting the bodies: Estimating the numbers and spatial variation of Australian reptiles, birds and mammals killed by two invasive mesopredators. Divers Distrib, 28, 976–991.

Thompson, S.D. (1987). Body size, duration of parental care, and the intrinsic rate of natural increase in eutherian and metatherian mammals. Oecologia, 71, 201–209.

Thomson, A.M., Calvin, K. V, Smith, S.J., Kyle, G.P., Volke, A., Patel, P., Delgado-Arias, S., Bond-Lamberty, B., Wise, M.A., Clarke, L.E. & Edmonds, J.A. (2011). RCP4.5: a pathway for stabilization of radiative forcing by 2100. Clim Change, 109, 77–94.

Tomlinson, S., Lomolino, M. V, Woinarski, J.C.Z., Murphy, B.P., Reed, E., Johnson, C.N., Legge, S., Helgen, K.M., Brown, S.C. & Fordham, D.A. (2023). Reconstructing mechanisms of extinctions to guide mammal conservation biogeography. J Biogeogr, 50, 1199–1212.

van der Vaart, E., Beaumont, M.A., Johnston, A.S.A. & Sibly, R.M. (2015). Calibration and evaluation of individual-based models using Approximate Bayesian Computation. Ecol Modell, 312, 182–190.

VanDerWal, J., Shoo, L.P., Johnson, C.N. & Williams, S.E. (2009). Abundance and the environmental niche: Environmental suitability estimated from niche models predicts the upper limit of local abundance. Am Nat, 174, 282–291.

Weeks, A.R., van Rooyen, A., Mitrovski, P., Heinze, D., Winnard, A. & Miller, A.D. (2013). A species in decline: genetic diversity and conservation of the Victorian eastern barred bandicoot, Perameles gunnii. Conservation Genetics, 14, 1243–1254.

Weeks, A.R., Sgrò, C.M., Young, A.G., Frankham, R., Mitchell, N.J., Miller, K.A., Byrne, M., Coates, D.J., Eldridge, M.D.B., Sunnucks, P., Breed, M.F., James, E.A. & Hoffmann, A.A. (2011). Assessing the benefits and risks of translocations in changing environments: a genetic perspective. Evol Appl, 4, 709–725.

Welch, J.J., Bininda-Emonds, O.R.P. & Bromham, L. (2008). Correlates of substitution rate variation in mammalian protein-coding sequences. BMC Evol Biol, 8, 53.

Westgate, M., Stevenson, M., Kellie, D. & Newman, P. (2024). galah: Atlas of Living Australia (ALA) Data and Resources in R.

Whitlock, M.C. & Bürger, R. (2009). Fixation of new mutations in small populations. In: Evolutionary conservation biology (eds. Ferrière, R., Dieckmann, U. & Couvet, D.). Cambridge University Press, Cambridge, UK, pp. 155–170.

Wiens, J.A., Stralberg, D., Jongsomjit, D., Howell, C.A. & Snyder, M.A. (2009). Niches, models, and climate change: assessing the assumptions and uncertainties. Proceedings of the National Academy of Sciences, 106, 19729–19736.

Willi, Y., Van Buskirk, J. & Hoffmann, A.A. (2006). Limits to the adaptive potential of small populations. Annu Rev Ecol Evol Syst, 37, 433–458.

Willi, Y., Kristensen, T.N., Sgrò, C.M., Weeks, A.R., Ørsted, M. & Hoffmann, A.A. (2022). Conservation genetics as a management tool: The five best-supported paradigms to assist the management of threatened species. Proceedings of the National Academy of Sciences, 119.

Wilson, B.A., Evans, M.J., Batson, W.G., Banks, S.C., Gordon, I.J., Fltecher, D.B., Wimpenny, C., Newport, J., Belton, E., Rypalski, A., Portas, T. & Manning, A.D. (2020). Adapting reintroduction tactics in successive trials increases the likelihood of establishment for an endangered carnivore in a fenced sanctuary. PLoS One, 15, e0234455.

Wilson, B.A., Evans, M.J., Gordon, I.J., Pierson, J.C., Brockett, B.M., Wimpenny, C., Batson, W.G., Newport, J. & Manning, A.D. (2023). Roadmap to recovery revealed through the reintroduction of an IUCN Red List species. Biodivers Conserv, 32, 227–248.

Winnard, A.L. & Coulson, G. (2008). Sixteen years of Eastern Barred Bandicoot *Perameles gunnii* reintroductions in Victoria: a review. Pacific Conservation Biology, 14, 34–53.

Withers, P.C., Cooper, C.E. & Larcombe, A.N. (2006). Environmental correlates of physiological variables in marsupials. Physiological and Biochemical Zoology, 79, 437– 453.

Woinarski, J.C.Z., Braby, M.F., Burbidge, A.A., Coates, D., Garnett, S.T., Fensham, R.J., Legge, S.M., McKenzie, N.L., Silcock, J.L. & Murphy, B.P. (2019). Reading the black book: The number, timing, distribution and causes of listed extinctions in Australia. Biol Conserv, 239, 108261.

Woinarski, J.C.Z., Burbidge, A.A. & Harrison, P.L. (2014). The action plan for Australian mammals 2012. CSIRO Publishing, Melbourne, Australia.

